# Quantitative assessment of bovine serum albumin proteins for blocking applications

**DOI:** 10.1101/869677

**Authors:** Gamaliel Junren Ma, Abdul Rahim Ferhan, Joshua A. Jackman, Nam-Joon Cho

**Affiliations:** School of Materials Science and Engineering, Nanyang Technological University, 50 Nanyang Avenue 639798, Singapore; School of Chemical Engineering, Sungkyunkwan University, Suwon 16419, Republic of Korea

## Abstract

Bovine serum albumin (BSA) is one of the most widely used protein reagents in the scientific community, especially for surface passivation (“blocking”) applications in various bioassays. Numerous BSA protein options are commercially available, however, there is scarce information about which ones are preferable for blocking applications. Herein, we conducted biophysical and bioassay measurements to quantitatively compare the conformational, adsorption, and blocking properties of BSA protein reagents that were obtained through six purification methods. Depending on the method, there were significant differences in the conformational and adsorption properties of BSA proteins, mainly due to the presence of fatty acid stabilizers. In turn, we discovered that fatty acid-free BSA proteins exhibit superior blocking performance to fatty acid-stabilized BSA proteins in surface- and nanoparticle-based bioassays. We critically discuss mechanistic factors behind these performance variations and our findings offer a practical framework to guide BSA selection for blocking applications.

---

A common element of bioassays is the need to minimize nonspecific binding of biomolecules in order to maximize assay specificity and sensitivity^1, 2^. To meet this objective, a wide range of reagents, ranging from crude materials such as milk proteins to synthetic nanomaterials such as finely tuned polymers, have been developed to coat target surfaces and to minimize nonspecific binding events—a function that is commonly referred to as “blocking”^3-7^. Among the various options, bovine serum albumin (BSA) is the most widely used protein reagent for blocking applications and has several practical advantages such as natural abundance, low cost, well-established purification methods, and broad availability. BSA is regarded as a gold-standard blocking reagent in numerous bioassays, including enzyme-linked immunosorbent assays (ELISA)^8, 9^, blots^10, 11^, immunohistochemistry^12, 13^, cell culture^14^, and biosensors^15, 16^. It is used as a coating to improve nanoparticle stability and biocompatibility^17, 18^, and is also commonly incorporated into buffer solutions to prevent the loss of precious materials such as nucleic acids, peptides, and enzymes^19, 20^.

Within this scope, an often-overlooked aspect of BSA as a blocking reagent is that there are many types of commercially available BSA that differ in terms of how the protein was isolated from bovine plasma, which is its principal natural source^21^. These processing differences relate to the choice of fractionation method and the use of chemical stabilizers, and such differences can cause significant variations in blocking performance in bioassays, even when comparing BSA reagents from the same company^22^. However, there is essentially no discussion in the scientific literature about what type(s) of BSA reagents are useful to pick for blocking applications, especially from a mechanistic perspective. Most blocking-related studies that investigate mechanistic details have only evaluated a single type of BSA and compare BSA adsorption and surface-induced conformational changes on different surfaces^23,24^ or investigate the effects of ionic strength^25^ or BSA pretreatment^26^. However, the comparison of BSA proteins obtained through different purification methods remains unexplored despite being the greatest source of variability in BSA reagent options on the market. Furthermore, there is limited reporting of the type of BSA reagent that is used in most scientific publications; the supplier company is sometimes listed, but the catalog number of the reagent is rarely provided. As such, BSA reagent selection for blocking applications largely remains a matter of trial-and-error and laboratory precedent, and establishing an evidence-backed practical framework to guide BSA reagent selection is an outstanding need.

Herein, the objective of our study was to investigate the conformational, adsorption, and blocking properties of commercially available BSA reagents that were obtained through six purification methods, and to develop a practical framework to guide BSA reagent selection for blocking applications. Our findings provide mechanistic insights to explain why certain types of BSA reagents are favorable over other options, and also outline key functional criteria to develop improved versions of BSA and other protein reagents for blocking applications.

## Results

### Evaluation strategy

BSA reagents are widely used to coat target surfaces in order to form surface passivation coatings, which minimize the nonspecific binding of interfering biomolecules that can affect bioassay readouts. The target surface can be a bulk surface such as a glass coverslip or polymeric membrane, or a nanostructured surface such as an inorganic nanoparticle. In practice, an experimentalist will incubate the target surface with an aqueous BSA solution, whereby protein molecules noncovalently adsorb and undergo a conformational change (“denature”) on the surface to form a thin film protein coating (**Fig. 1a**).

**Figure 1.**
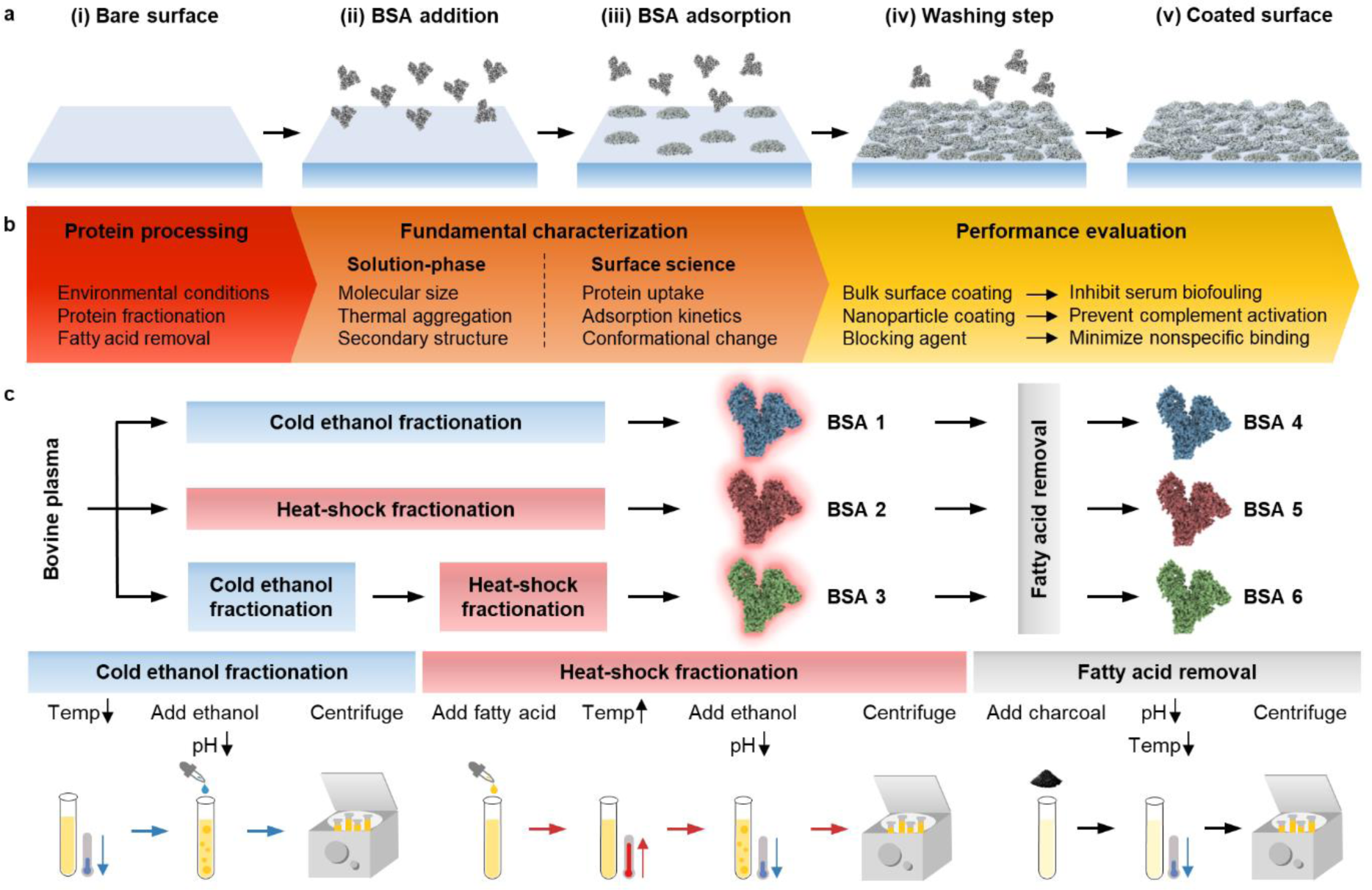
Evaluation of BSA protein reagents for blocking applications. **a**, Overview of basic steps in BSA blocking protocols. (i) A target surface is identified to coat with BSA for surface passivation (“blocking”). (ii) BSA proteins in aqueous solution are incubated with the target surface. (iii) BSA proteins adsorb and denature on the target surface. (iv) A washing step is performed to remove weakly bound BSA protein molecules. (v) A firmly attached monolayer of adsorbed BSA protein molecules is formed on the surface. **b**, Overall experimental strategy to analyze BSA reagent properties. A series of BSA protein reagents, which were purified by different methods, were selected and their conformational, adsorption, and blocking properties were evaluated. A series of solution-phase and surface-sensitive biophysical measurement approaches were employed along with surface- and nanoparticle-based bioassays. **c**, Outline of purification routes to isolate BSA protein molecules from bovine plasma. Fatty acid-containing BSA proteins (referred to as “fatted”) were prepared by (1) cold ethanol fractionation, (2) heat-shock fractionation, and (3) cold ethanol fractionation followed by heat-shock fractionation and these reagents are designated as BSA 1, 2, 3, respectively. Fatty acid-free versions (referred to as “defatted”) of BSA 1, 2, 3 were also prepared and are designated as BSA 4, 5, 6, respectively.

Based on this coating process, our evaluation strategy focused on characterizing the conformational, adsorption, and blocking properties of six commercial BSA reagents that were purified by different methods (**Fig. 1b**). We employed a wide range of solution-phase and surface-sensitive biophysical measurement tools to characterize the BSA reagents along with surface- and nanoparticle-based bioassay application testing. This approach allowed us to not only directly compare the functional performance of different BSA reagents in blocking applications, but to also obtain key mechanistic insights into how subtle differences in the conformational properties of proteins – as influenced by the processes involved in each purification method – affect protein adsorption, surface-induced denaturation, and resulting blocking performance.

We selected six commercial BSA reagents that were purified through three different fractionation routes, without or with a fatty acid removal step (**Fig. 1c** and **Supplementary Table 1**). The fractionation step is necessary in order to separate BSA from other serum components and there are two main options: cold ethanol fractionation and heat-shock fractionation. Briefly, cold ethanol fractionation involves reducing the sample temperature to approximately −5 °C followed by the addition of ethanol and adjusting the solution pH and ionic strength in order to isolate BSA protein^27-29^. By contrast, heat-shock fractionation involves heating the sample (typically to ∼60 °C or slightly higher) in order to isolate BSA protein and is conducted in the presence of a fatty acid stabilizer (caprylic acid)^30-32^. These two fractionation processes can be completed alone or sequentially (cold ethanol, then heat-shock).

After the fractionation step(s) is completed, purified BSA reagents are obtained. Notably, at this stage, all of the BSA reagents still contain fatty acids that were either naturally bound or were added as a stabilizer^33^. We tested fatty acid-containing BSA reagents (referred to as “fatted”) that were prepared by (1) cold ethanol fractionation, (2) heat-shock fractionation, and (3) cold ethanol fractionation followed by heat-shock fractionation and these reagents were designated as BSA 1, 2, 3, respectively. To remove fatty acids, an extra processing step is required, which involves the addition of activated charcoal followed by adjusting solution pH and temperature^34^. We also tested the fatty acid-free versions (referred to as “defatted”) of BSA 1, 2, 3 and they were designated as BSA 4, 5, 6, respectively. In total, six different BSA reagents were tested consisting of three fatty acid-containing (1-3) and three fatty acid-free (4-6) versions.

### Thermal stability of fatted and defatted BSA proteins

Like other globular proteins, BSA folds in aqueous solution in order to minimize its conformational free energy. The folding pathway is dictated by the net balance of stabilizing and destabilizing forces, and the resulting folded structure is typically only a few kcal/mol lower in free energy than the unfolded amino acid chain^35^. Thus, subtle differences in the purification method – such as the fractionation route, thermal history, and fatty acid content – can have important consequences on the conformational free energy of BSA protein molecules. Such effects can manifest themselves by influencing the thermal stability of proteins in the solution phase as well as by modulating adsorption and surface-induced denaturation profiles on a target surface.

As a first step, we investigated the thermal stability of the different BSA reagents by conducting dynamic light scattering (DLS) and circular dichroism (CD) spectroscopy experiments (**Fig. 2a**). The experiments were conducted in temperature-controlled measurement chambers, starting at 25 °C and the temperature was gradually raised in 5 °C increments. The DLS technique measures the size distribution of protein molecules in bulk solution and could detect the onset of temperature-induced protein oligomerization, as indicated by a marked increase in the average hydrodynamic diameter. The size increase occurs when thermally denatured protein monomers re-assemble into oligomeric structures^36^. A higher onset temperature indicates greater thermal stability and vice-versa. On the other hand, the CD spectroscopy technique identifies the secondary structure characteristics of protein molecules in bulk solution, and we tracked the decrease in α-helical character with increasing temperature due to protein unfolding.

**Figure 2.**
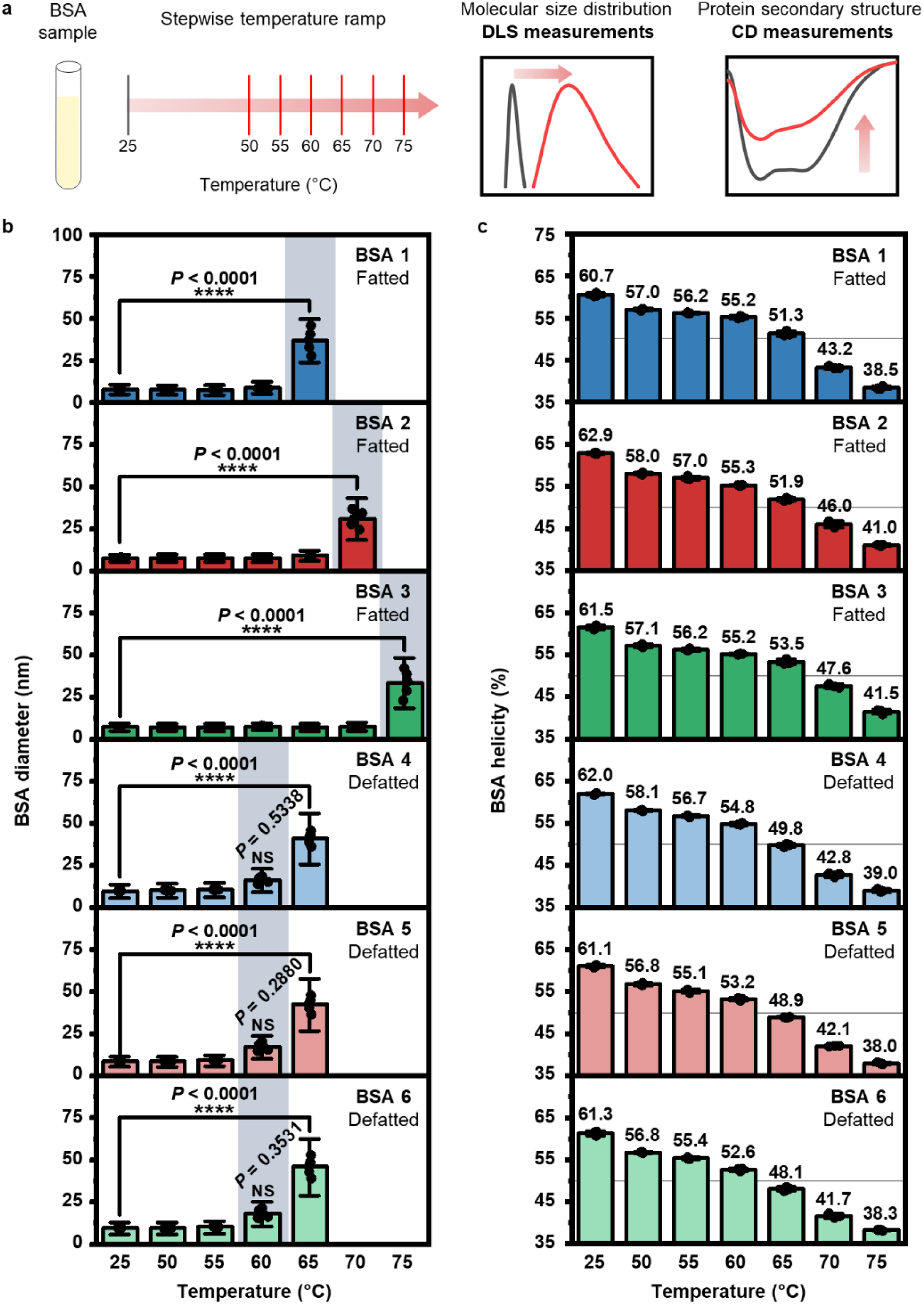
Temperature profiling of BSA conformational stability in solution. **a**, Schematic illustration of DLS and CD spectroscopy measurement strategies. Experiments were conducted in temperature-controlled measurement chambers and measurements were recorded in 5 °C increments (in increasing order). **b**, Hydrodynamic diameter of BSA protein molecules as a function of temperature, as measured in DLS experiments. Data are reported as mean ± standard deviation (s.d.), where s.d. is defined as the full-width-at-half-maximum (FWHM)/2.355 [*n*=5 technical replicates, one-way analysis of variance (ANOVA) with Dunnett’s multiple comparisons test (versus data at 25 °C)]. Dots represent individual data points. **c**, Fractional percentage of α-helicity in BSA protein molecules as a function of temperature, as measured in CD spectroscopy experiments. Mean values are presented on top of each column. Values were computed from molar residue ellipticity data and data are reported as mean ± s.d. (*n*=3 technical replicates). Dots represent individual data points. The grey horizontal line in each graph corresponds to 50% helicity.

Across the temperature range of 25 to 55 °C, the DLS data showed that all six BSA proteins have ∼8-nm diameter, which agrees well with the expected size of BSA monomers^25, 26, 37, 38^ (**Fig. 2b** and **Supplementary Fig. 1**). At 60 °C, a marked increase in the sizes of defatted BSA proteins (4-6) was detected, with average diameters around 20 nm that indicated the onset of protein oligomerization. By contrast, the sizes of fatted BSA proteins (1-3) remained stable and did not increase at 60 °C. At 65 °C, the sizes of the defatted BSA proteins (4-6) increased to around 45 nm diameter and the onset of protein oligomerization occurred for one of the fatted BSA proteins (1), with an average diameter around 35 nm. By contrast, the onset temperatures for the other fatted BSA proteins (2) and (3) were 70 and 75 °C, respectively. By extending the incubation time to 2 hours and fixing the temperature at 60 °C, it was verified that defatted BSA proteins are prone to more rapid and extensive oligomerization while fatted BSA proteins are more thermally stable (**Supplementary Fig. 2**).

Thermally induced oligomerization of BSA molecules is triggered by irreversible conformational changes in the protein structure once there is a critical degree of protein unfolding and corresponding loss of α-helical character. The CD spectroscopy data showed that all six BSA proteins exhibited around 60-63% α-helical character at 25 °C, followed by a progressive decrease in α-helical character with increasing temperature^25, 26, 39, 40^ (**Fig. 2c** and **Supplementary Figs. 3**,**4**). Both reversible conformational changes (below the onset temperature of oligomerization) and irreversible conformational changes (at and above the onset temperature) contribute to protein unfolding so we compared the helicity data at 65 °C. All three defatted BSA proteins sample (4-6) had helical percentages below 50% on average, while all three fatted BSA protein samples (1-3) had helical percentages above 51% on average (**Supplementary Table 2**). More pronounced differences in helicity were observed at 70 °C, and mirroring the DLS data, the findings also support that, among the fatted BSA samples, BSA protein (1) had the lowest helical percentage and hence, lowest thermal stability. The relatively low thermal stability of BSA protein (1) is consistent with its processing by cold ethanol fractionation alone, and thus only natural fatty acids are bound in the absence of additional fatty acid stabilizer (as used in heat-shock fractionation). Collectively, the DLS and CD spectroscopy findings demonstrate that fatted BSA samples have greater thermal stability than defatted BSA samples, supporting that the conformational free energy of fatted BSA samples is lower than defatted BSA samples.

### Adsorption behavior of fatted and defatted BSA proteins

These findings led us to investigate the adsorption properties of BSA proteins on hydrophilic silica surfaces (**Fig. 3a)**. When a protein with higher conformational energy adsorbs onto a solid surface, it is expected to undergo greater unfolding and spreading on the surface due to entropic gains while other factors such as protein-surface and protein-protein interactions also affect the outcome^41, 42^. We conducted quartz crystal microbalance-dissipation (QCM-D) experiments in order to track the adsorption kinetics and corresponding mass and viscoelastic properties of adsorbed BSA protein molecules. The QCM-D technique measures real-time changes in the resonance frequency (ΔF) and energy dissipation (ΔD) properties of a silica-coated sensor surface as a function of time, which provide insight into the hydrodynamically-coupled mass and viscoelastic properties of adsorbed BSA protein |ΔF_max_| shift molecules on the silica surface, respectively. A larger indicates greater protein adsorption, while a larger |ΔF/ΔD| shift reflects greater rigidity (less viscoelasticity) within the adsorbed protein layer.

**Figure 3.**
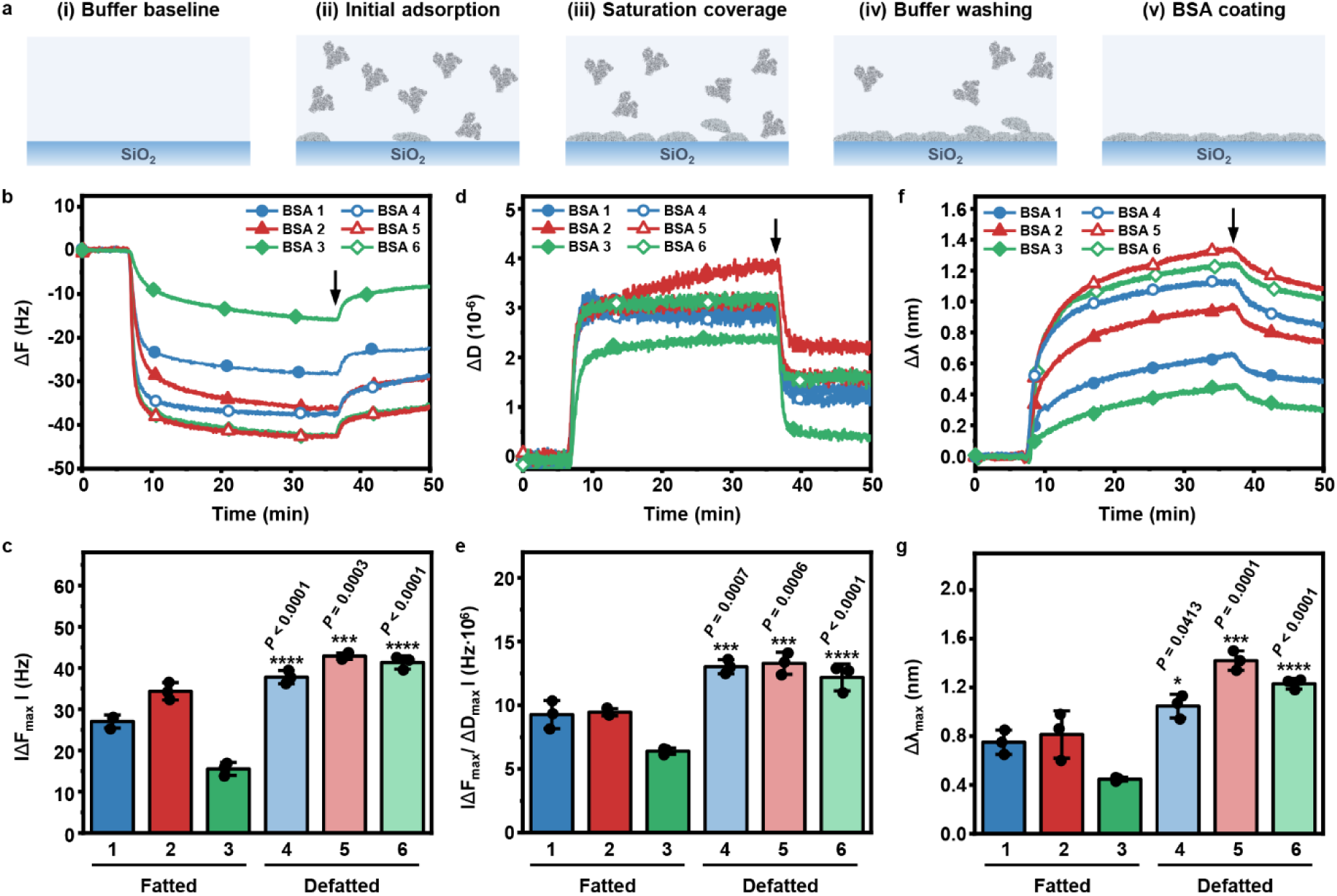
Real-time monitoring of BSA protein adsorption onto silica surfaces. **a**, Overview of experimental protocol for testing BSA reagents. (i) A baseline signal is established in aqueous buffer solution (ii) 100 μM BSA in equivalent buffer solution is added (initiated at *t*=5 min). (iii) BSA adsorption continues until approaching saturation uptake. (iv) A buffer washing step is performed (initiated at *t*=35 min and indicated by an arrow in panels **b, d, f**), and (v) an adsorbed BSA protein layer is formed. **b-e**, QCM-D experiments were conducted to measure frequency (ΔF) and energy dissipation (ΔD) signals related to the protein adsorption process. **b**, Time-resolved QCM-D ΔF shifts and **c**, corresponding |ΔF_max_| shifts at saturation. **d**, Time-resolved QCM-D ΔD shifts. **e**, |ΔF_max_/ΔD_max_| ratios obtained from saturation data in panels **b** and **d. f-g**, LSPR experiments were conducted to measure Δλ_max_ signals related to the protein adsorption process. **f**, Time-resolved LSPR wavelength shifts (Δλ) and **g**, corresponding Δλ_max_ shifts at saturation. Data in **c, e, g** are reported as mean ± s.d. (*n*=3 biological replicates, one-way ANOVA with Tukey’s multiple comparisons test). *P* values are reported for defatted BSA proteins (versus fatted BSA proteins from the same fractionation method). Dots represent individual data points.

All six BSA proteins adsorbed onto the silica surface and the QCM-D frequency signals indicated monotonic adsorption kinetics (**Fig. 3b**). Most adsorbed protein molecules were |ΔF_max_| bound irreversibly as judged by a buffer washing step. In general, there were larger shifts of around 35 to 45 Hz for defatted BSA proteins (4-6), indicating greater adsorption (**Fig. 3c**). By contrast, there was less adsorption of fatted BSA proteins (1-3), as indicated by |ΔF_max_| shifts around 15 to 35 Hz. The corresponding QCM-D energy dissipation signals also indicated monotonic protein adsorption and some degree of viscoelastic character within the adsorbed protein layer (**Fig. 3d**). Viscoelastic model fitting estimated that the protein adlayers have effective thicknesses on the order of 3-6 nm, which corresponds to a single layer of adsorbed protein molecules (**Supplementary Fig. 5**). After a buffer washing step, the remaining bound protein molecules exhibited a more rigid arrangement. Further analysis of the |ΔF_max_/ΔD_max_| ratio revealed that adsorbed protein layers from defatted BSA proteins (4-6) were more tightly bound to the silica surface, which is consistent with lower conformational stability (in solution) and greater surface-induced denaturation (**Fig. 3e** and **Supplementary Fig. 6**). On the other hand, fatted BSA proteins (1-3) exhibited greater viscoelastic character indicative of less surface-induced denaturation.

To corroborate the QCM-D results, we also performed localized surface plasmon resonance (LSPR) experiments to measure BSA protein adsorption onto silica-coated surfaces. While the QCM-D technique is sensitive to adsorbed protein mass and hydrodynamically-coupled solvent mass, the LSPR technique detects the wavelength shift (Δλ) that is associated with changes in local refractive index near the sensor surface and the measurement signal is only sensitive to the adsorbed protein mass (not coupled solvent)^38^. In LSPR experiments, the Δλ shift is tracked as a function of time, whereby a larger Δλ shift indicates greater protein adsorption. For all six BSA proteins, a positive Δλ shift occurred and monotonic adsorption kinetics were observed (**Fig. 3f**). After a buffer washing step, most protein molecules remained bound. Importantly, the Δλ_max_ shifts showed that the defatted BSA proteins exhibited greater adsorption uptake than the fatted BSA proteins (∼1.2 ± 0.2 nm vs. ∼0.6 ± 0.2 nm), which agrees well with the QCM-D data (**Fig. 3g**). Thus, defatted BSA proteins exhibited higher adsorption uptake and tighter adlayer packing, whereas fatted BSA proteins had lower adsorption uptake and weaker adlayer packing.

### Adsorption-related protein conformational changes

To further compare the extent of surface-induced protein denaturation for each BSA protein, we analyzed the LSPR measurement data by computing the initial rate of adsorption uptake, which is denoted as (dΔλ/dt)_max_ (**Supplementary Fig. 7**). The LSPR technique is a highly surface-sensitive measurement approach and it is possible to relate the initial adsorption rate to the relative extent of surface-induced protein denaturation, whereby a larger initial rate is indicative of greater denaturation^40, 43^. Consequently, the LSPR data showed that defatted BSA proteins undergo more extensive surface-induced denaturation, as indicated by initial adsorption rates around 0.6-0.7 nm min^−1^ (**Fig. 4a**). On the other hand, the initial adsorption rates for fatted BSA proteins were only around 0.2-0.3 nm min^−1^, which indicate less surface-induced denaturation.

**Figure 4.**
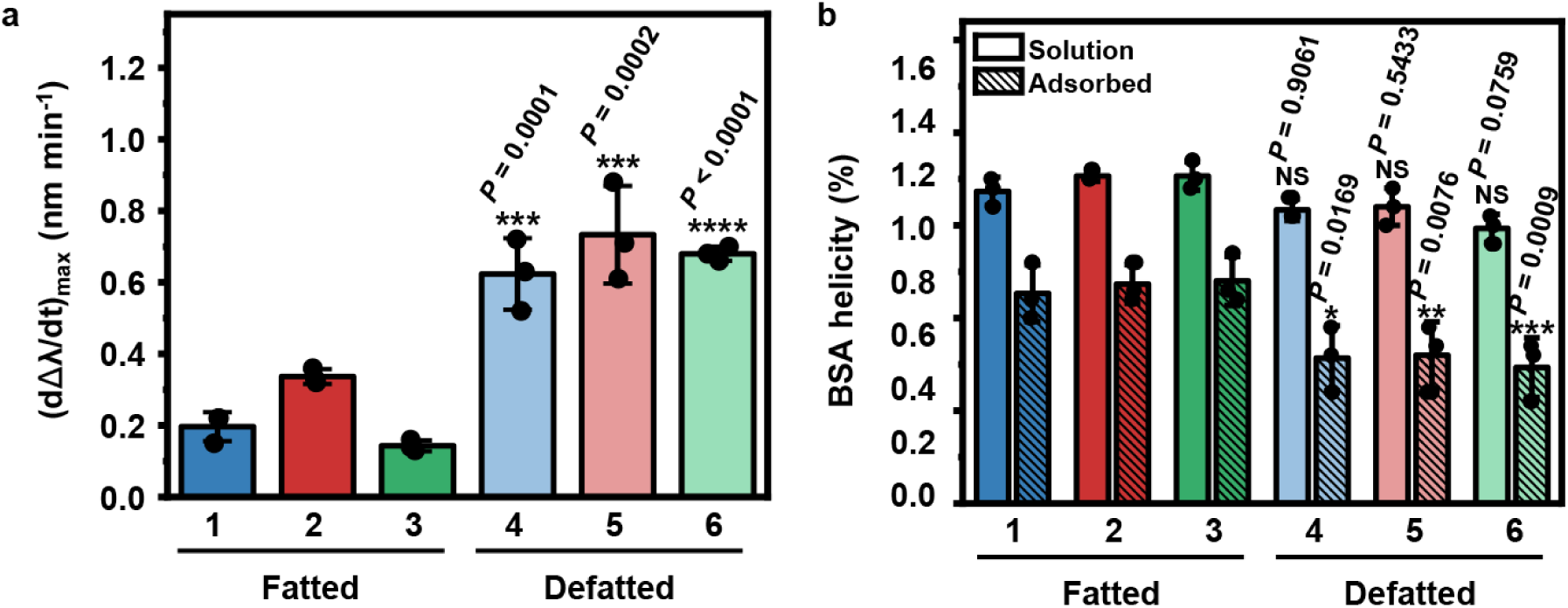
Quantitative evaluation of adsorption-related conformational changes in BSA protein structure. **a**, Maximum rate of change in the LSPR wavelength shift(dΔλ/dt)_max_ during the initial adsorption stage. Values are computed from data in Fig. 3f. Data are reported as mean ± s.d. (*n*=3 biological replicates, one-way ANOVA with Tukey’s multiple comparisons test). **b**, Fractional percentage of α-helicity in BSA protein molecules in solution and in the adsorbed state, as determined in ATR-FTIR spectroscopy experiments. Data are reported as mean ± s.d. (*n*=3 biological replicates, two-way ANOVA with Tukey’s multiple comparisons test). *P* values are reported for defatted BSA proteins (versus fatted BSA proteins from the same fractionation method) in **a**, and are separately reported for defatted BSA proteins in solution (versus fatted BSA proteins in solution from the same fractionation method) and for defatted BSA proteins in the adsorbed state (versus fatted BSA proteins in the adsorbed state from the same fractionation method) in **b**. Dots represent individual data points.

In addition, attenuated total reflection-Fourier transform infrared (ATR-FTIR) spectroscopy measurements were performed to measure the change in BSA protein structure due to adsorption-related conformational changes. In general, in the solution phase, fatted BSA molecules had α-helicities around 63% while defatted ones had α-helicities around 60%, which agree with literature values^44, 45^ (**Fig. 4b, Supplementary Fig. 8**, and **Supplementary Table 3**). Upon adsorption, fatted BSA molecules underwent surface-induced denaturation and the fractional helicities decreased to around 53%, indicating a net helical loss of ∼10%. By contrast, upon adsorption, defatted BSA molecules underwent greater surface-induced denaturation and the resulting fractional helicities were around 45%, which corresponds to a net helical loss of ∼15%. Thus, multiple lines of experimental data support that defatted BSA molecules exhibit greater surface-induced denaturation than fatted BSA molecules. This finding is consistent with the lower conformational stability exhibited by defatted BSA molecules in solution, and supports that decreased conformational stability of a solution-phase protein translates into more pronounced denaturation in the adsorbed state.

### Surface coating application performance

As numerous applications of BSA protein molecules are linked to their role as surface coatings, we next investigated the application performance of the six BSA samples in surface- and nanoparticle-based bioassays. We first measured the blocking efficiency of adsorbed BSA coatings to inhibit serum biofouling on silica surfaces (**Fig. 5a** and **Supplementary Fig. 9**). A bare silica surface was first coated with BSA protein molecules before incubation of the BSA-coated surface in 100% fetal bovine serum (FBS) for 30 min, followed by a washing step. The resulting amount of serum biofouling on the BSA-coated silica surface was evaluated by the QCM-D technique. The blocking efficiency of the BSA coating was calculated based on the degree to which serum biofouling was inhibited. It was determined that defatted BSA proteins outperformed fatted BSA proteins, and defatted BSA proteins (5) and (6) had around 90% blocking efficiency (**Fig. 5b**). On the other hand, fatted BSA proteins (1) and (3) demonstrated less than 40% blocking efficiency. These findings are consistent with greater adsorption uptake and higher packing density of adsorbed, defatted BSA protein molecules. Similar performance results were also obtained when fatted and defatted BSA proteins were tested in Western blot experiments involving a nitrocellulose membrane surface (**Supplementary Figs. 10-12** and **Supplementary Note 1**).

**Figure 5.**
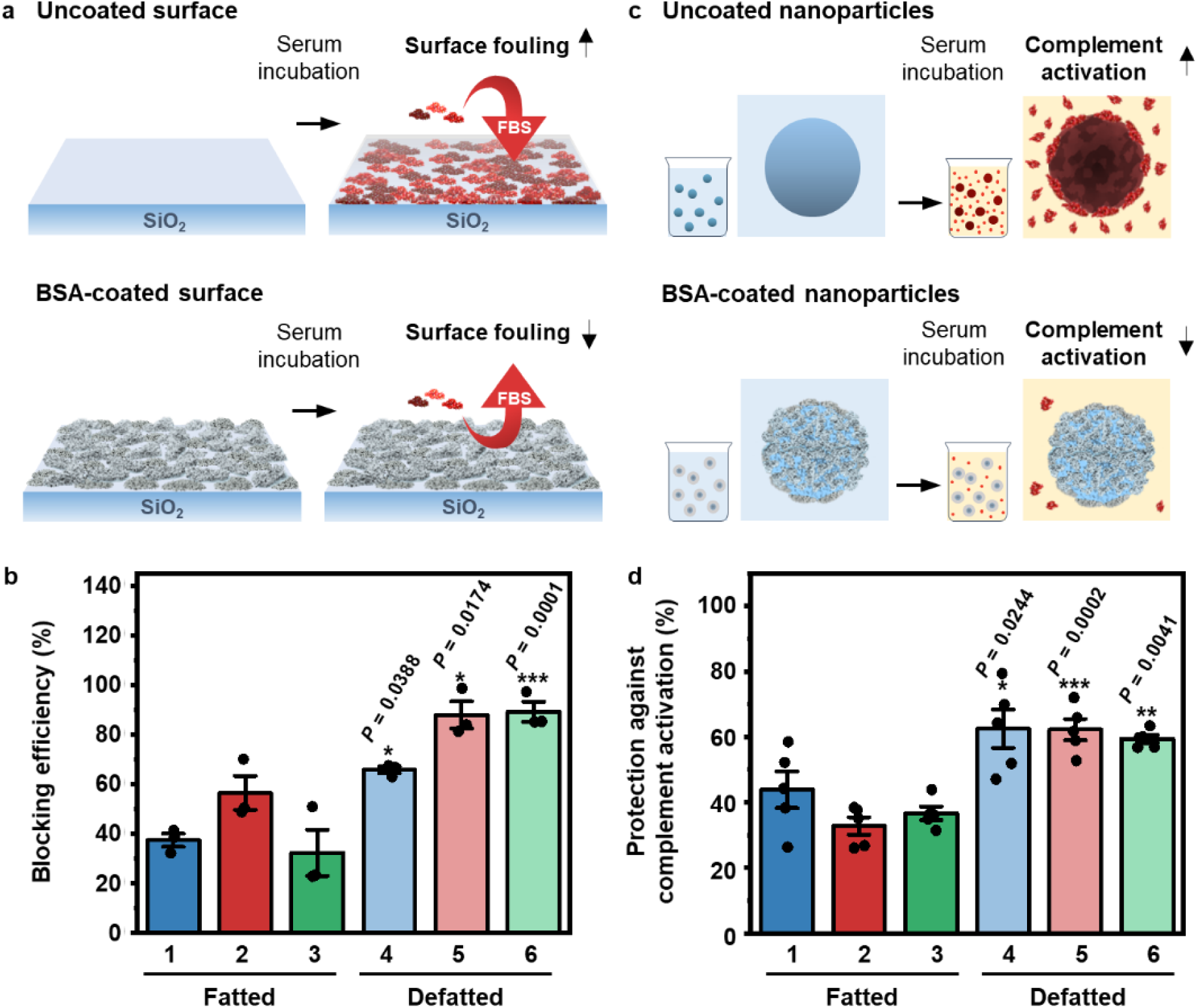
Blocking performance of BSA protein reagents in surface- and nanoparticle-based bioassays. **a**, Schematic illustration of serum biofouling assay on a silica surface. Pristine and BSA-coated silica surfaces were incubated in 100% fetal bovine serum (FBS) followed by a buffer washing step, and the amount of serum biofouling on the surfaces was determined by QCM-D measurements. The blocking efficiency of the BSA coatings was calculated relative to the uncoated surface. **b**, Blocking efficiencies of BSA coatings to inhibit serum biofouling on silica surfaces (*n*=3 biological replicates). **c**, Schematic illustration of nanoparticle-triggered complement activation assay. Pristine and BSA-coated silica nanoparticles were incubated in 100% normal human serum (NHS), followed by ELISA measurements to determine the extent of nanoparticle-triggered complement activation, as indicated by the magnitude of sC5b-9 production. Protection efficiency of BSA coatings was calculated relative to uncoated nanoparticles. **d**, Protection efficiencies of BSA coatings to inhibit nanoparticle-induced complement activation (*n*=5 biological replicates). Data in **b** and **d** are reported as mean ± s.e.m. (one-way ANOVA with Tukey’s multiple comparisons test). *P* values are reported for defatted BSA proteins (versus fatted BSA proteins from the same fractionation method). Dots represent individual data points.

We also tested the blocking performance of BSA coatings on silica nanoparticles to inhibit nanoparticle-induced complement activation, which is an acute immune response that can occur in human serum (**Fig. 5c**)^46-49^. Using an enzyme-linked immunosorbent assay (ELISA), we measured the degree of protection which BSA coatings conferred against nanoparticle-induced complement activation, as judged by the amount of SC5b-9 protein biomarker in solution (**Fig. 5d** and **Supplementary Fig. 13**). Greater SC5b-9 protein concentrations indicate more extensive complement activation and conversely nanoparticles with more protective coatings would trigger lower SC5b-9 protein concentration levels compared to bare nanoparticles. Experimentally, it was determined that defatted BSA coatings inhibited more than 60% of SC5b-9 generation, whereas the fatted BSA coatings exhibited 40% or less protection efficiency. This finding supports that defatted BSA coatings are superior to fatted BSA coatings to protect against nanoparticle-induced complement activation. Thus, across the surface- and nanoparticle-based assays, defatted BSA proteins exhibited better functional performance than fatted BSA proteins.

## Discussion

Our experimental findings demonstrate that defatted BSA proteins exhibit distinct conformational and adsorption properties from fatted BSA proteins, and these differences lead to significant variations in blocking performance. Overall, the data point to the central role of fatty acid molecules in affecting BSA structure and function. To verify the effect of fatty acids, we also doped defatted BSA with caprylic acid, thereby converting the defatted protein into a fatted protein. Caprylic acid addition led to markedly enhanced conformational stability along with decreased adsorption uptake and denaturation and consequently poorer blocking performance (**Supplementary Figs. 14-23, Supplementary Tables 4-5**, and **Supplementary Note 2**). Mechanistically, the enhanced conformational stability of fatted BSA proteins can be understood through the insertion of caprylic acid molecules (via hydrophobic tails) into hydrophobic pockets on the molecular surface of BSA proteins, which confers a net stabilizing effect on protein structure^50, 51^.

A less obvious but critically important effect of caprylic acid on BSA structure relates to the protein adsorption process itself (**Fig. 6**). When a protein adsorbs onto a surface, the protein will typically undergo surface-induced denaturation and the extent of protein denaturation depends on a combination of factors, including the protein’s intrinsic conformational stability and the magnitude of protein-surface interactions^42^. More denaturation causes greater protein spreading on the surface, which means that the contact surface area per adsorbing protein molecule is larger. As such, it is generally understood that greater spreading of individual protein molecules results in a smaller total number of adsorbed protein molecules on a target surface, irrespective of whether the greater spreading is due to conformational stability (see, *e.g.*, Ref. 52) or protein-surface interactions (see, *e.g.*, Ref. 53). From this perspective, it is possible to rationalize how changing ionic strength affects BSA protein adsorption and spreading for a single type of BSA^25^.

**Figure 6.**
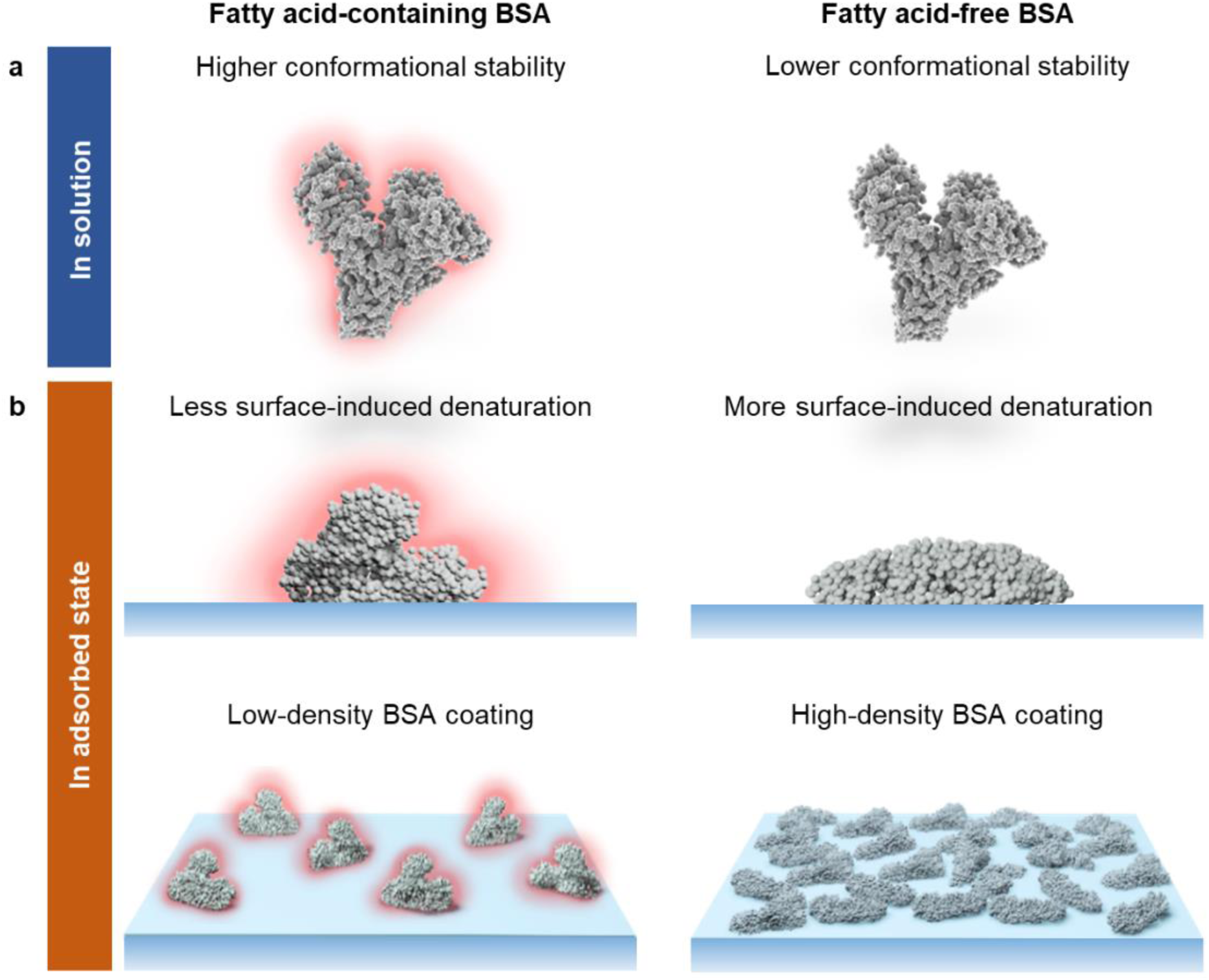
Effect of fatty acids on BSA conformational and adsorption properties and implications for blocking applications. **a**, Schematic illustration of fatted and defatted BSA protein molecules in solution. The presence of fatty acids leads to higher conformational stability in solution. **b**, Schematic illustration of fatted and defatted BSA protein molecules in the adsorbed state. Higher conformational stability of fatted BSA leads to less adsorption-induced denaturation, greater protein-protein repulsion, and thus less well-packed protein coatings. On the other hand, the lower conformational stability of defatted BSA leads to greater adsorption-induced denaturation, less protein-protein repulsion, and thus more well-packed protein coatings. As a result, defatted BSA proteins form more effective surface passivation coatings that are more useful for blocking applications.

However, the situation is more nuanced when comparing defatted and fatted BSA proteins because caprylic acid molecules not only enhance conformational stability but the protruding carboxylic acid headgroups of caprylic acid molecules also increase the negative surface charge character of BSA protein molecules. Fatted BSA proteins have greater conformational stability and thus undergo less surface-induced denaturation in the adsorbed state. Nevertheless, while fatted BSA proteins have a smaller contact surface area per molecule, the maximum surface density of adsorbed protein molecules in this case is limited by repulsive electrostatic interactions between nearest-neighbor protein molecules.

On the other hand, defatted BSA proteins have lower conformational stability and denature to a greater extent in the adsorbed state. Consequently, defatted BSA proteins undergo greater spreading and have a larger contact surface area per adsorbed molecule. However, the maximum surface density of adsorbed proteins in the defatted BSA case is still higher than in the fatted BSA case because there are less repulsive protein-protein interactions between adsorbed, defatted BSA molecules. Hence, defatted BSA proteins are useful to form passivation coatings for two main reasons. First, they have lower conformational stability that permits greater denaturation and spreading in the adsorbed state. Second, the absence of fatty acid molecules in defatted BSA proteins permits more favorable protein-protein interactions that support the formation of close-packed, adsorbed protein layers on target surfaces.

Taken together, our findings support that it is preferable to use fatty acid-free BSA protein reagents for blocking applications across surface- and nanoparticle-based bioassays. In general, fatty acid-free BSA proteins outperformed fatty acid-stabilized BSA proteins largely independent of the fractionation route that was used in the purification stage. Mechanistically, we show that defatted BSA proteins are superior blocking reagents because they have lower conformational stability in solution, which translates into greater adsorption and surface-induced denaturation as well as tighter adlayer packing that collectively lead to superior passivation coatings. Depending on the assay specifics, additional consideration of BSA purity level (*i.e.*, residual levels of one or more specific plasma components) can also be warranted, while fatty acid-free status is the main selection criteria to ensure optimal blocking properties. Looking forward, our analytical approach outlines a series of important functional properties – solution-phase conformational stability, adsorption uptake, surface-induced denaturation, and molecular packing within the protein coating – that are important to take into account when evaluating protein-based reagents for blocking applications and can also guide the development of improved versions of BSA and other protein-based blocking reagents alone and as part of multi-component buffer solutions.

## Acknowledgements

The authors thank Mr. Tun Naw Sut for technical assistance with QCM-D experiments. This work was supported by the National Research Foundation of Singapore through a Competitive Research Programme grant (NRF-CRP10-2012-07) and a Proof-of-Concept grant (NRF2015NRF-POC0001-19), and by the Creative Materials Discovery Program through the National Research Foundation of Korea (NRF) funded by the Ministry of Science, ICT, and Future Planning (NRF-2016M3D1A1024098).

## Author Contributions

G.J.M, A.R.F, J.A.J., and N.-J.C. planned the studies. G.J.M. and A.R.F. conducted experiments. G.J.M., A.R.F., J.A.J., and N.-J.C. interpreted the results. J.A.J. and N.-J.C. obtained funding. G.J.M., A.R.F., and J.A.J. wrote the first draft of the manuscript. All authors reviewed, edited and approved the paper.

## Competing Interests

The authors declare no competing interests.

## Methods

### Bovine serum albumin proteins

The six types of bovine serum albumin (BSA) used in this study were selected based on three different fractionation methods without or with a fatty acid removal step, and were procured from Sigma-Aldrich (St. Louis, MO, USA). Lyophilized powders of fatty acid-containing BSA proteins purified by cold ethanol fractionation, heat-shock fractionation, or cold ethanol followed by heat-shock fractionation were selected and have catalog nos. A2153, A3059, and A7638, respectively. These three fatted BSA proteins were labeled BSA 1, 2, 3, respectively. The corresponding fatty acid-free versions were also obtained and have catalog nos. A6003, A7030 and A0281, respectively. These three defatted BSA proteins were labeled BSA 4, 5, 6, respectively.

### Reagents

Sodium dodecyl sulfate (SDS, catalog no. L4390), sodium chloride (NaCl, catalog no. 746398), sodium hydroxide (catalog no. S5881) and octanoic acid (caprylic acid, catalog no. C2875) were also purchased from Sigma-Aldrich while Tris(hydroxymethyl)aminomethane (Tris, catalog no. 0497) was purchased from Amresco (Solon, OH, USA). Ethanol (95%) was purchased from Aik Moh (Singapore), hydrochloric acid (HCl, catalog no. 100317) was purchased from Merck (Burlington, MA, USA) and fetal bovine serum (FBS, catalog no. SV30160.03, lot no. RC35960) was purchased from HyClone Laboratories (Logan, UT, USA) and stored at −20 °C until experiment. Normal human serum (catalog no. NHS, lot no. 38) was obtained from Complement Technology (Tyler, TX, USA) and stored at −80 °C until experiment. 30% Acrylamide/Bis Solution 29:1 (catalog no. 1610156), ammonium persulfate (catalog no. 1610700), tetramethylethylenediamine (TEMED, catalog no. 1610800), 4× Laemmli sample buffer (catalog no. 1610747), 2-mercaptoethanol (catalog no. 1610710), 10× Tris buffered saline (TBS, catalog no. 1706435), 10× Tris/glycine buffer (catalog no. 1610734), 10× Tris/glycine/SDS buffer (catalog no. 1610732), Tween 20 (catalog no. 1706531), nitrocellulose membranes (catalog no. 1620112), Precision Plus Protein Standards (catalog no. 1610375), Clarity Max Western enhanced chemiluminescent (ECL) substrate (catalog no. 1705062), and horseradish peroxidase (HRP)-conjugated goat anti-mouse immunoglobulin G (IgG) antibody (catalog no. 1721011, batch no. 64109318) were all purchased from Bio-Rad Laboratories (Hercules, CA, USA). Complement C3b monoclonal antibody (catalog no. MA1-40155, lot no. SH2428445) was purchased from Thermo Fisher Scientific (Waltham, MA, USA). Methanol (catalog no. M/4000/17) was obtained from Fisher Scientific (Loughborough, UK) and 100-nm diameter silica nanoparticles (catalog no. SISN100) were obtained from nanoComposix (San Diego, CA, USA).

### Sample preparation

An aqueous buffer solution of 10 mM Tris, 150 mM NaCl, and pH 7.5 was prepared with Milli-Q-treated water (resistivity of >18.2 MΩ.cm at 25 °C) and filtered through a 0.2 μm polyethersulfone (PES) membrane filter (Thermo Fisher Scientific, catalog no. 595-4520). BSA solutions were prepared by dissolving lyophilized BSA powder in this Tris buffer and then filtering through a 0.2 μm syringe filter (catalog no. PN-4612; Pall Corporation, Port Washington, NY, USA). The molar concentrations of BSA proteins in aqueous buffer solution were determined by ultraviolet (UV) light absorption measurements at 280 nm and with a molar extinction coefficient value of 43,824 M^−1^ cm^−1^. Where applicable, the addition of caprylic acid to BSA samples was conducted by first dissolving caprylic acid and sodium hydroxide into the Tris buffer in order to make a 50 mM caprylic acid solution at pH 7.5. The caprylic acid solution was then added to appropriate amounts of BSA solution in order to yield caprylic acid-doped BSA sample with a molar ratio of 10:1 caprylic acid:BSA. In addition, the Western blot running buffer was prepared by diluting 10× Tris/glycine/SDS buffer with water to 1× concentration, and the transfer buffer was prepared by diluting the 10× Tris/glycine buffer with water and methanol to 1× concentration and adding 20% (vol/vol) of methanol. The BSA-containing blocking solutions for Western blot experiments were prepared by first diluting 10× TBS with water, followed by the addition of Tween 20 to make a 1× TBS solution with 0.1% (vol/vol) Tween 20 (TBST). BSA was then dissolved in TBST to make a 3% (wt/vol) BSA blocking solution.

### Dynamic light scattering

Dynamic light scattering (DLS) was employed to measure the average hydrodynamic diameter and polydispersity of BSA protein molecules in solution using a particle size analyzer (ZetaPALS, Brookhaven Instruments, Holtsville, NY, USA) that is equipped with a 658.0 nm monochromatic laser. Measurements were taken at a 90° scattering angle and the BIC Particle Sizing software (v5.27; Brookhaven Instruments) was used to analyze the intensity autocorrelation function in order to obtain the intensity-weighted Gaussian size distribution. The temperature in the measurement chamber was controlled with a feedback loop and measurements of a protein sample were first recorded at 25 °C, followed by heating and increasing the temperature from 50 °C to 75 °C in 5 °C increments. After each temperature step increase, the measurement chamber was equilibrated for 5 min before the DLS measurement was performed. In select time-dependent measurements, the temperature in the measurement chamber was first raised from 50 °C to 55 °C before maintaining a constant temperature at 60 °C and measuring protein size every 10 min for 200 min. All reported values were obtained from 5 technical replicates.

### Circular dichroism spectroscopy

Circular dichroism (CD) spectroscopy experiments were conducted using an AVIV Model 420 CD spectrometer with the AVIV CDS software package (v3.36 MX) (AVIV Biomedical, Lakewood, NJ, USA). CD spectroscopy measurements were conducted using 400 μL solutions of 2.5 μM BSA samples in a 1 mm path length cuvette with a PTFE stopper (catalog no. 110-QS; Hellma, Müllheim, Germany). The measurements were recorded with a 1 nm spectral bandwidth, 0.5 nm step size, and an averaging time of 0.1 s. Elevated temperature experiments were conducted by increasing the temperature in the measurement chamber in 5 °C increments from 50 to 75 °C and recording the spectra at every temperature point after a 5 min equilibration time. All resulting spectra were processed by subtracting a background spectra of equivalent solution conditions without BSA protein and the data were presented in mean residue molar ellipticity ([θ]) units based on the following equation: 

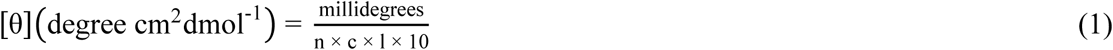

where *n* is the number of amino acid residues, *c* is the protein molar concentration, and *l* is the cuvette path length in cm. All CD spectra were smoothed by using the Savitzky-Golay^54^ smoothing function with a smoothing window of 20 points and a polynomial order of 2 in the OriginPro 2019b (v9.6.5.169) software package (OriginLab, Northampton, MA, USA). The α-helical percentage of each BSA protein was calculated from the [θ] data at 222 nm ([θ]_222_) based on the following equation^55^: 

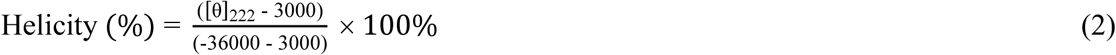

### Quartz crystal microbalance-dissipation monitoring

Quartz crystal microbalance-dissipation (QCM-D) measurements were conducted using a QSense E4 instrument (Biolin Scientific AB, Stockholm, Sweden). A silica-coated AT-cut quartz crystal sensor chip with a fundamental frequency of 5 MHz (QSX 303, Biolin Scientific) was used to characterize protein adsorption onto silica surfaces. The temperature was set to 25 °C for all experiments, and the measurement operation was controlled by the QSoft 401 (v2.5.13.664) (Biolin Scientific) software package. Prior to each experiment, the sensor chips were sequentially rinsed with 1% (wt/vol) aqueous SDS solution, water, and ethanol, and then dried under a gentle stream of nitrogen gas, followed by treatment with oxygen plasma (PDC-002, Harrick Plasma, Ithaca, NY, USA) for 3 min. A peristaltic pump was used to inject liquid samples into the measurement chamber at a volumetric flow rate of 100 μL min^−1^. A stable baseline signal was first established in Tris buffer solution before a 100 μM BSA protein was introduced into the measurement chamber for 30 min, followed by a buffer washing step. The resonance frequency (ΔF) and energy dissipation (ΔD) shifts were recorded in real-time at multiple odd overtones, as previously described^56^, and the normalized data at the fifth overtone are reported. The QCM-D data were also analyzed by the Voigt-Voinova model^57^ using the QTools (v3.0.15.553) (Biolin Scientific) software package in order to estimate the effective thickness of the adsorbed protein layer formed after buffer rinsing. For this analysis, the adsorbed protein layer was assumed to be a homogenous, single-layer of protein molecules with uniform thickness and with a uniform effective density of 1300 kg m^−3^. The density and viscosity of the bulk aqueous buffer solution were fixed at 1000 kg m^−3^ and 0.001 Pa s^−1^, respectively.

The QCM-D technique was also employed to determine the blocking efficiency of BSA coatings against serum biofouling. For these experiments, 100 μM BSA solution was first added to a bare silica surface under continuous flow for 60 min, followed by a 40 min buffer washing step. Undiluted fetal bovine serum (FBS) was then introduced under continuous flow for 80 min, followed by a final buffer washing step for 30 min. The blocking efficiency percentage was calculated by comparing the absolute difference between the frequency shift due to BSA adsorption alone (after first buffer washing step) and the subsequent frequency shift due to FBS biofouling (after second buffer washing step), which is denoted as ΔF_FBS-BSA_. A control experiment without BSA coating was also conducted in order to measure the absolute frequency shift due to FBS biofouling alone (after buffer washing step), which is denoted as ΔF_Control_. For each BSA coating, the blocking efficiency percentage was calculated by the following equation: 

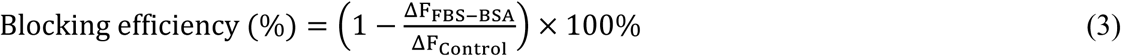

### Localized surface plasmon resonance

Ensemble-averaged localized surface plasmon resonance (LSPR) measurements were conducted in optical transmission mode using an Insplorion XNano instrument (Insplorion AB, Gothenburg, Sweden), as previously described^58^. A white light beam illuminating a circular area of ∼4 mm^2^ is transmitted through a silica-coated sensor chip (Insplorion) consisting of well-separated gold nanodisk arrays (∼8% surface coverage) deposited by hole-mask colloidal lithography^59^ on a transparent glass substrate. The nanodisks had an average height and diameter of 20 and 120 nm. Respectively .and the surface of the sensor chip was conformally coated with a silica film (∼10 nm thick). The light was then passed through a quartz glass window and collected by a spectrophotometer. Prior to each experiment, the sensor chip was sequentially rinsed with 1% (wt/vol) aqueous SDS solution, water and ethanol, then dried under a gentle stream of nitrogen gas. The chip was then treated with oxygen plasma for 2.5 min. The sensor chip was then loaded into the measurement chamber and liquid samples were introduced by a peristaltic pump at a volumetric flow rate of 100 μL min^−1^. A stable baseline signal was established in Tris buffer solution before 100 μM BSA was introduced into the measurement chamber under continuous flow for 30 min, followed by a buffer washing step. The Insplorer software package (Insplorion AB) was used to record the LSPR extinction spectra with a time resolution of 1 Hz and the centroid (peak) position (λ) in the extinction spectrum at each time point was calculated by using a high-order polynomial fitting^60^. Thus, it was possible to determine the time-resolved Δλ shift due to protein adsorption and the time-derivative plot of the Δλ shift was also calculated using the OriginPro 2019b software package.

### Attenuated total reflection-Fourier transform infrared spectroscopy

Attenuated total reflection-Fourier transform infrared (ATR-FTIR) spectroscopy experiments were conducted using a Bruker Vertex 70 FTIR spectrometer with a liquid-nitrogen-cooled mercury cadmium telluride (MCT) photodetector (Bruker, Baden-Württemberg, Germany) that was equipped with a MIRacle ATR accessory containing a three-reflection ZnSe ATR crystal (PIKE Technologies, Fitchburg, WI, USA). A 30 μL aliquot of 100 μM BSA solution was pipetted onto the ATR crystal to form a droplet, and an absorbance spectrum was immediately recorded to characterize the secondary structure of solution-phase BSA proteins. The BSA was then incubated with the ATR crystal surface at room temperature for 30 min, followed by a series of ten buffer washing steps to remove weakly adsorbed proteins. During each washing step, 30 μL of extra buffer solution (without protein) was added to the sample droplet, followed by aspiration of a 30 μL total volume from the droplet. After the washing steps were completed, then a second absorbance spectrum was recorded to characterize the secondary structure of BSA proteins in the adsorbed state. During measurements, the sample compartment was purged continuously with nitrogen gas to minimize ambient moisture. The OPUS 6.5 software package (Bruker) was used to collect the ATR-FTIR absorbance spectra at a 4 cm^−1^ resolution and averaged over 128 scans. All recorded spectra were normalized by subtracting background spectra from buffer and water vapor alone in order to remove water and ambient water vapor absorbance artefacts, respectively.

The relative contribution of different secondary structure elements was determined for all samples by curve fitting the amide I bands of the absorbance spectra (1720 – 1580 cm^−1^) to Gaussian components based on the peaks identified in the second derivative plot of the absorbance spectra and from literature references ^44, 61-64^. All curve fitting analyses were conducted using the Peak Analyzer function in the OriginPro 2019b software package. The absorbance spectrum obtained from experiment was assumed to be a linear combination of multiple constituent Gaussian peaks, each centered around a given wavenumber value and representing a protein secondary structure element. The secondary structure percentage values were determined by calculating the area under the curve-fitted peaks that had been assigned to each secondary structure element as a fraction of the total area under the fitted curve. Peaks centered at 1682-1678 cm^−1^ were assigned to β-turn structures, 1656-1654 cm^−1^ to α-helix, 1649-1648 cm^−1^ to random coil, 1639-1624 cm^−1^ to extended chains, and 1618-1612 cm^−1^ to intermolecular β-sheets.

### Sodium dodecyl sulfate-polyacrylamide gel electrophoresis and Western blot

10% polyacrylamide resolving gels at pH 8.8 and 5% stacking gels at pH 6.8 with fifteen sample loading wells per gel were prepared before experiments. For sample preparation, normal human serum (NHS) was thawed and then immediately diluted to 1% (vol/vol) in Tris buffer, followed by incubation at 37 °C for 30 min. A 180 μL aliquot of diluted NHS was then mixed with 54 μL of 4× Laemmli buffer and 6 μL of 2-mercaptoethanol. The sample mixture was then heated to 95 °C for 5 min for complete denaturation of serum proteins, before cooling back to room temperature. A 10 μL aliquot of protein standards was loaded into the first lane of the polyacrylamide gel and the next two lanes were each loaded with 10 μL of the sample mixture; this sequence was repeated five times in total to fill all lanes within the gel. Sodium dodecyl sulfate-polyacrylamide gel electrophoresis (SDS-PAGE) was then conducted at 50 V for 15 min, followed by 100 V for 90 min in running buffer.

The protein bands in the gel were then transferred to a nitrocellulose membrane at 300 mA for 2 h in transfer buffer, and ice packs were used to keep the transfer buffer cool during the transfer process. After the transfer step, the membranes were cut into strips, each one containing the three lanes of protein standards (lane 1) and serum samples (lanes 2-3). Blocking solutions were prepared using fatted or defatted BSA [3% (wt/vol) BSA in TBST; BSA 1 or 5] and each strip was incubated with 5 mL of the appropriate BSA blocking solution at room temperature for 1 h. The membranes were then incubated with complement C3b monoclonal antibody as the primary antibody (1:500 dilution in blocking solution) overnight. Each membrane was then washed with TBST for 10 min. After three washing steps, the membranes were incubated with HRP-conjugated goat anti-mouse IgG secondary antibody (1:2000 dilution in blocking solution) for 1 h, followed by four subsequent washing steps with TBST.

The membranes were then incubated with Clarity Max Western ECL substrate for 5 min in order to enhance chemiluminescence properties. All incubation and washing steps were conducted on a rocking platform. The protein bands were subsequently imaged using an Amersham Imager 600 instrument and its accompanying software (v0.9.8) (GE Healthcare, Chicago, IL, USA) with a 1 s exposure time. The Fiji/ImageJ software package^65^ was then used to quantify the intensity of the noise bands that were located near the 250, 75, and 50 kDa molecular weight markers. The digital image of the blots was first set to 8-bit greyscale, and then an intensity profile from each selected lane was plotted using the Analyze Gels function. The peaks from non-specific bands of interest were identified and a baseline was created by drawing a straight line between the two minima points on either side of each peak of interest. The Wand tool was then used to determine the area bounded by the peak and the baseline. These values were defined as the intensity values for each selected non-specific band.

### Enzyme-linked immunosorbent assay

The MicroVue SC5b-9 Plus Enzyme Immunoassay kit (catalog no. A020; Quidel, San Diego, CA, USA) was used for enzyme-linked immunosorbent assay (ELISA) experiments. BSA-coated silica nanoparticle samples were prepared by mixing equal volumes of the appropriate 1 mg mL^−1^ BSA solution with 2 mg mL^− 1^ silica nanoparticles in Tris buffer. The mixtures were then incubated at 37 °C for 2 h, followed by centrifugation at 16,000×*g* for 30 min. The supernatants were removed and the nanoparticles were resuspended with fresh Tris buffer. Another round of centrifugation and resuspension was conducted to yield 1 mg mL^−1^ BSA-coated silica nanoparticle samples. A 10 μL aliquot of each BSA-coated silica nanoparticle sample was then mixed with a 40 μL aliquot of freshly thawed normal human serum (NHS). A 50 μL aliquot of NHS by itself was used as the negative control (lowest level of complement activation) and 10 μL of 1 mg mL^−1^ uncoated silica nanoparticles plus 40 μL NHS was used as the positive control (highest level of complement activation). All test samples were incubated at 37 °C for 30 min.

The assay was then performed using the 96-well plates that were provided in the ELISA kit according to the manufacturer’s instructions. All serum samples, including controls, were diluted 40-times by using the provided sample diluent solution before addition to the wells. Standard solutions were added without dilution. The absorbance values from each well were determined by an Infinite Pro 200 plate reader (Tecan, Männedorf, Switzerland). Standard curves were calculated by correlating the known concentrations of the standard solutions to the respective absorbance values. The SC5b-9 protein concentration in each sample was then computed. The degree of protection afforded by each BSA coating was calculated by treating the SC5b-9 protein concentrations from the negative and positive control samples as 100% and 0% protection, respectively.

### Statistical analysis

Statistical analyses were conducted using the GraphPad Prism (v8.0.1) software package from GraphPad Software (La Jolla, CA, USA). One-way or two-way analysis of variance (ANOVA) with the appropriate multiple comparisons test and the unpaired t-test were used to compute the statistical significance of experimental data as appropriate. All statistical analysis involved two-tailed tests. Unpaired t-tests results are reported as *P* values while multiple comparisons test results are reported as multiplicity-adjusted *P* values. *P* < 0.05, *P* < 0.01, *P* < 0.001 and *P* < 0.0001 indicate the levels of statistical significance. Additional information can be found in Supplementary Tables 6-15.

### Data Availability

All data supporting the findings of this study are available within this Article and its Supplementary Information files, and from the corresponding authors upon reasonable request.

## References

1. Nakanishi, K., Sakiyama, T. & Imamura, K. On the adsorption of proteins on solid surfaces, a common but very complicated phenomenon. J. Biosci. Bioeng. 91, 233–244 (2001).

2. Gray, J.J. The interaction of proteins with solid surfaces. Curr. Opin. Struct. Biol. 14, 110–115 (2004).

3. Nuzzo, R.G. Biomaterials: stable antifouling surfaces. Nat. Mater. 2, 207 (2003).

4. Miller, R.T., Kubier, P., Reynolds, B., Henry, T. & Turnbow, H. Blocking of endogenous avidin-binding activity in immunohistochemistry: the use of skim milk as an economical and effective substitute for commercial biotin solutions. Appl. Immunohisto. M. M. 7, 63–65 (1999).

5. Kirschner, C.M. & Brennan, A.B. Bio-inspired antifouling strategies. Annu. Rev. Mater. Res. 42, 211–229 (2012).

6. Wei, Q. et al. Protein interactions with polymer coatings and biomaterials. Angew. Chem. Int. Edit. 53, 8004–8031 (2014).

7. Lowe, S., O’Brien-Simpson, N.M. & Connal, L.A. Antibiofouling polymer interfaces: poly (ethylene glycol) and other promising candidates. Polym. Chem. 6, 198–212 (2015).

8. Avrameas, S. & Guilbert, B. Enzyme-immunoassay for the measurement of antigens using peroxidase conjugates. Biochimie 54, 837–842 (1972).

9. Jitsukawa, T., Nakajima, S., Sugawara, I. & Watanabe, H. Increased coating efficiency of antigens and preservation of original antigenic structure after coating in ELISA. J. Immunol. Methods 116, 251–257 (1989).

10. Towbin, H., Staehelin, T. & Gordon, J. Electrophoretic transfer of proteins from polyacrylamide gels to nitrocellulose sheets: procedure and some applications. Proc. Natl. Acad. Sci. 76, 4350–4354 (1979).

11. Mahmood, T. & Yang, P.-C. Western blot: technique, theory, and trouble shooting. N. Am. J. Med. Sci. 4, 429 (2012).

12. Ramos-Vara, J. Technical aspects of immunohistochemistry. Vet. Pathol. 42, 405–426 (2005).

13. Buchwalow, I.B. & Böcker, W. in Immunohistochemistry: Basics and Methods 41–46 (Springer Berlin Heidelberg, Berlin, Heidelberg; 2010).

14. Francis, G.L. Albumin and mammalian cell culture: implications for biotechnology applications. Cytotechnology 62, 1–16 (2010).

15. Špačková, B., Wrobel, P., Bocková, M. & Homola, J. Optical biosensors based on plasmonic nanostructures: a review. Proc. IEEE 104, 2380–2408 (2016).

16. Ronkainen, N.J., Halsall, H.B. & Heineman, W.R. Electrochemical biosensors. Chem. Soc. Rev. 39, 1747–1763 (2010).

17. Brewer, S.H., Glomm, W.R., Johnson, M.C., Knag, M.K. & Franzen, S. Probing BSA binding to citrate-coated gold nanoparticles and surfaces. Langmuir 21, 9303–9307 (2005).

18. Mariam, J., Sivakami, S. & Dongre, P.M. Albumin corona on nanoparticles–a strategic approach in drug delivery. Drug Deliv. 23, 2668–2676 (2016).

19. Kreader, C.A. Relief of amplification inhibition in PCR with bovine serum albumin or T4 gene 32 protein. Appl. Environ. Microbiol. 62, 1102–1106 (1996).

20. Goebel-Stengel, M., Stengel, A., Taché, Y. & Reeve Jr, J.R. The importance of using the optimal plasticware and glassware in studies involving peptides. Anal. Biochem. 414, 38–46 (2011).

21. Mannuzza, F.J. & Montalto, J.G. Is bovine albumin too complex to be just a commodity? Bioprocess Int. (2010).

22. Xiao, Y. & Isaacs, S.N. Enzyme-linked immunosorbent assay (ELISA) and blocking with bovine serum albumin (BSA)—not all BSAs are alike. J. immunol. Methods 384, 148–151 (2012).

23. Sweryda-Krawiec, B., Devaraj, H., Jacob, G. & Hickman, J.J. A new interpretation of serum albumin surface passivation. Langmuir 20, 2054–2056 (2004).

24. Jeyachandran, Y., Mielczarski, J., Mielczarski, E. & Rai, B. Efficiency of blocking of non-specific interaction of different proteins by BSA adsorbed on hydrophobic and hydrophilic surfaces. J. Colloid Interface Sci. 341, 136–142 (2010).

25. Park, J.H. et al. Controlling adsorption and passivation properties of bovine serum albumin on silica surfaces by ionic strength modulation and cross-linking. Phys. Chem. Chem. Phys. 19, 8854–8865 (2017).

26. Park, J.H. et al. Temperature-induced denaturation of BSA protein molecules for improved surface passivation coatings. ACS Appl. Mater. Interfaces 10, 32047–32057 (2018).

27. Cohn, E.J. et al. Preparation and properties of serum and plasma proteins. IV. A system for the separation into fractions of the protein and lipoprotein components of biological tissues and fluids. J. Am. Chem. Soc. 68, 459–475 (1946).

28. Kistler, P. & Nitschmann, H. Large scale production of human plasma fractions: eight years experience with the alcohol fractionation procedure of Nitschmann, Kistler and Lergier. Vox Sang. 7, 414–424 (1962).

29. Hao, Y.I. A simple method for the preparation of human serum albumin. Vox Sang. 36, 313–320 (1979).

30. Hoch, H. & Chanutin, A. Albumin from heated human plasma. I. Preparation and electrophoretic properties. Arch. Biochem. Biophys. 51, 271–276 (1954).

31. Reid, A.F. (Google Patents, 1955).

32. Schneider, W., Lefevre, H., Fiedler, H. & McCarty, L.J. An alternative method of large scale plasma fractionation for the lsolation of serum albumin. Blut 30, 121–134 (1975).

33. Spector, A.A. Fatty acid binding to plasma albumin. J. Lipid Res. 16, 165–179 (1975).

34. Chen, R.F. Removal of fatty acids from serum albumin by charcoal treatment. J. Biol. Chem. 242, 173–181 (1967).

35. Dobson, C.M., Šali, A. & Karplus, M. Protein folding: a perspective from theory and experiment. Angew. Chem. Int. Edit. 37, 868–893 (1998).

36. Militello, V. et al. Aggregation kinetics of bovine serum albumin studied by FTIR spectroscopy and light scattering. Biophys. Chem. 107, 175–187 (2004).

37. Jachimska, B., Wasilewska, M. & Adamczyk, Z. Characterization of globular protein solutions by dynamic light scattering, electrophoretic mobility, and viscosity measurements. Langmuir 24, 6866–6872 (2008).

38. Jackman, J.A., Ferhan, A.R. & Cho, N.-J. Nanoplasmonic sensors for biointerfacial science. Chem. Soc. Rev. 46, 3615–3660 (2017).

39. Moriyama, Y. et al. Secondary structural change of bovine serum albumin in thermal denaturation up to 130 C and protective effect of sodium dodecyl sulfate on the change. J. Phys. Chem. B 112, 16585–16589 (2008).

40. Jackman, J.A. et al. Indirect nanoplasmonic sensing platform for monitoring temperature-dependent protein adsorption. Anal. Chem. 89, 12976–12983 (2017).

41. Norde, W. in Macromol. Symp., Vol. 103 5–18 (Wiley Online Library, 1996).

42. Rabe, M., Verdes, D. & Seeger, S. Understanding protein adsorption phenomena at solid surfaces. Adv. Colloid Interface Sci. 162, 87–106 (2011).

43. Jackman, J.A. et al. Nanoplasmonic ruler to measure lipid vesicle deformation. Chem. Commun. 52, 76–79 (2016).

44. Givens, B.E., Xu, Z., Fiegel, J. & Grassian, V.H. Bovine serum albumin adsorption on SiO_2_ and TiO_2_ nanoparticle surfaces at circumneutral and acidic pH: A tale of two nano-bio surface interactions. J. Colloid Interface Sci. 493, 334–341 (2017).

45. Kossovsky, N. et al. Secondary structure of albumin acquired rapidly by modified conventional ATR-FTIR is comparable to CD spectral data. J. Colloid interface Sci. 166, 350–355 (1994).

46. Ricklin, D., Hajishengallis, G., Yang, K. & Lambris, J.D. Complement: a key system for immune surveillance and homeostasis. Nat. Immunol. 11, 785 (2010).

47. Szebeni, J. Complement activation-related pseudoallergy: a stress reaction in blood triggered by nanomedicines and biologicals. Mol. Immunol. 61, 163–173 (2014).

48. Belling, J.N. et al. Stealth immune properties of graphene oxide enabled by surface-bound complement Factor H. ACS Nano 10, 10161–10172 (2016).

49. Fülöp, T. et al. Complement activation in vitro and reactogenicity of low-molecular weight dextran-coated SPIONs in the pig CARPA model: Correlation with physicochemical features and clinical information. J. Control. Release 270, 268–274 (2018).

50. Curry, S., Mandelkow, H., Brick, P. & Franks, N. Crystal structure of human serum albumin complexed with fatty acid reveals an asymmetric distribution of binding sites. Nat. Struct. Mol. Biol. 5, 827 (1998).

51. Curry, S. in Advances in Molecular and Cell Biology, Vol. 33 29–46 (2003).

52. Karlsson, M., Ekeroth, J., Elwing, H. & Carlsson, U. Reduction of irreversible protein adsorption on solid surfaces by protein engineering for increased stability. J. Biol. Chem. 280, 25558–25564 (2005).

53. Ramsden, J. & Prenosil, J. Effect of ionic strength on protein adsorption kinetics. J. Phys. Chem. 98, 5376–5381 (1994).

## References

54. Savitzky, A. & Golay, M.J. Smoothing and differentiation of data by simplified least squares procedures. Anal. Chem. 36, 1627–1639 (1964).

55. Morrisett, J.D., David, J.S., Pownall, H.J. & Gotto Jr, A.M. Interaction of an apolipoprotein (apoLP-alanine) with phosphatidylcholine. Biochemistry 12, 1290–1299 (1973).

56. Cho, N.-J., Frank, C.W., Kasemo, B. & Höök, F. Quartz crystal microbalance with dissipation monitoring of supported lipid bilayers on various substrates. Nat. Protoc. 5, 1096 (2010).

57. Voinova, M.V., Rodahl, M., Jonson, M. & Kasemo, B. Viscoelastic acoustic response of layered polymer films at fluid-solid interfaces: continuum mechanics approach. Phys. Scr. 59, 391 (1999).

58. Jackman, J.A., Zhdanov, V.P. & Cho, N.-J. Nanoplasmonic biosensing for soft matter adsorption: kinetics of lipid vesicle attachment and shape deformation. Langmuir 30, 9494–9503 (2014).

59. Fredriksson, H. et al. Hole–mask colloidal lithography. Adv. Mater. 19, 4297–4302 (2007).

60. Dahlin, A.B., Tegenfeldt, J.O. & Höök, F. Improving the instrumental resolution of sensors based on localized surface plasmon resonance. Anal. Chem. 78, 4416–4423 (2006).

61. Murayama, K. & Tomida, M. Heat-induced secondary structure and conformation change of bovine serum albumin investigated by Fourier transform infrared spectroscopy. Biochemistry 43, 11526–11532 (2004).

62. Kong, J. & Yu, S. Fourier transform infrared spectroscopic analysis of protein secondary structures. Acta Biochim. Biophys. Sin. 39, 549–559 (2007).

63. Barth, A. Infrared spectroscopy of proteins. BBA-Bioenergetics 1767, 1073–1101 (2007).

64. Srour, B., Bruechert, S., Andrade, S.L. & Hellwig, P. in Membrane Protein Structure and Function Characterization 195–203 (Springer, 2017).

65. Schindelin, J. et al. Fiji: an open-source platform for biological-image analysis. Nat. Methods 9, 676 (2012).

