## Supporting Information for "Quantitative assessment of bovine serum albumin proteins for blocking applications"

### Supplementary Note 1

### Quantification of nonspecific band intensities in Western blot experiments

In Western blot experiments, the blocking step prior to antibody incubation is a critical part of the protocol to minimize nonspecific adsorption of primary and/or secondary antibodies onto the blot membrane, which can lead to false signals and poor blot resolution. To this end, BSA is the gold-standard blocking reagent although there is no discussion in the scientific literature about the blocking performance of fatted versus defatted BSA proteins.

Therefore, we conducted Western blot experiments and used normal human serum (NHS) as our protein sample against a C3b monoclonal primary antibody, which produces two specific bands corresponding to C3b (104 kDa) and iC3b (40 kDa) proteins. While these two bands are clearly visible in all Western blots, we also identified a few additional nonspecific bands near the 250, 75, and 50 kDa molecular weight markers. Our experimental results demonstrated that defatted BSA significantly decreased the intensity of these nonspecific bands, as compared to fatted BSA (**Supplementary Fig. 10**). This observation was confirmed by quantitative evaluation of the chemiluminescence intensity values of the nonspecific bands, as determined by Fiji/ImageJ software[1]. The method used to quantify the band intensities by Fiji/ImageJ software is illustrated in **Supplementary Fig. 11** and the details are described in the Methods section.

### Supplementary Note 2

#### Addition of caprylic acid to defatted and fatted BSA proteins

To verify that the observed differences in the conformational and adsorption properties of fatted and defatted BSA proteins are due to fatty acids, we treated BSA protein 5 with caprylic acid supplementation in order to convert defatted BSA 5 into fatted CA-BSA 5 (10:1 molar ratio of caprylic acid to BSA 5 protein). Caprylic acid is the most widely used fatty acid in the BSA fractionation process in order to stabilize protein molecules against temperature-induced denaturation[2-4]. The corresponding data are presented in **Supplementary Figs. 14-22** and **Supplementary Tables 4** and **5**. Overall, the data support that fatty acid-containing CA-BSA 5 is more stable than fatty acid-free BSA 5, with respect to thermal denaturation and adsorption-related surface denaturation. This finding confirms the important role of fatty acids in modulating the conformational and adsorption properties of BSA proteins.

While defatted BSA proteins performed quite similarly in all tested assays, we also noticed that the fatted BSA proteins showed some degree of variation in conformational stability depending on the assay. Among the fatted BSA proteins, BSA 3 typically showed the highest levels of conformational stability, as reflected across solution-phase and surface-sensitive biophysical measurements. This led us to suspect that the fatted BSA proteins had different degrees of "fatting" since conformational stability is related to fatty acid-protein ratio[5] (higher fatty acid content yields greater stability). Therefore, we supplemented the fatted BSA proteins 1-3 with additional caprylic acid (10:1 molar ratio of caprylic acid to BSA proteins) in order to see the effects on QCM-D adsorption kinetics (**Supplementary Fig. 23**). Negligible changes in BSA 3 adsorption behavior were observed without or with caprylic acid doping, thereby confirming that the as-supplied BSA 3 was fully fatted. By contrast, there tended to be slight differences in the adsorption behavior of BSA 1 and 2 without and with caprylic acid doping, especially with respect to adlayer rigidity. Overall, the caprylic acid supplementation experiments verified that the fatted BSA proteins 1-3 all have high fatty acid contents.

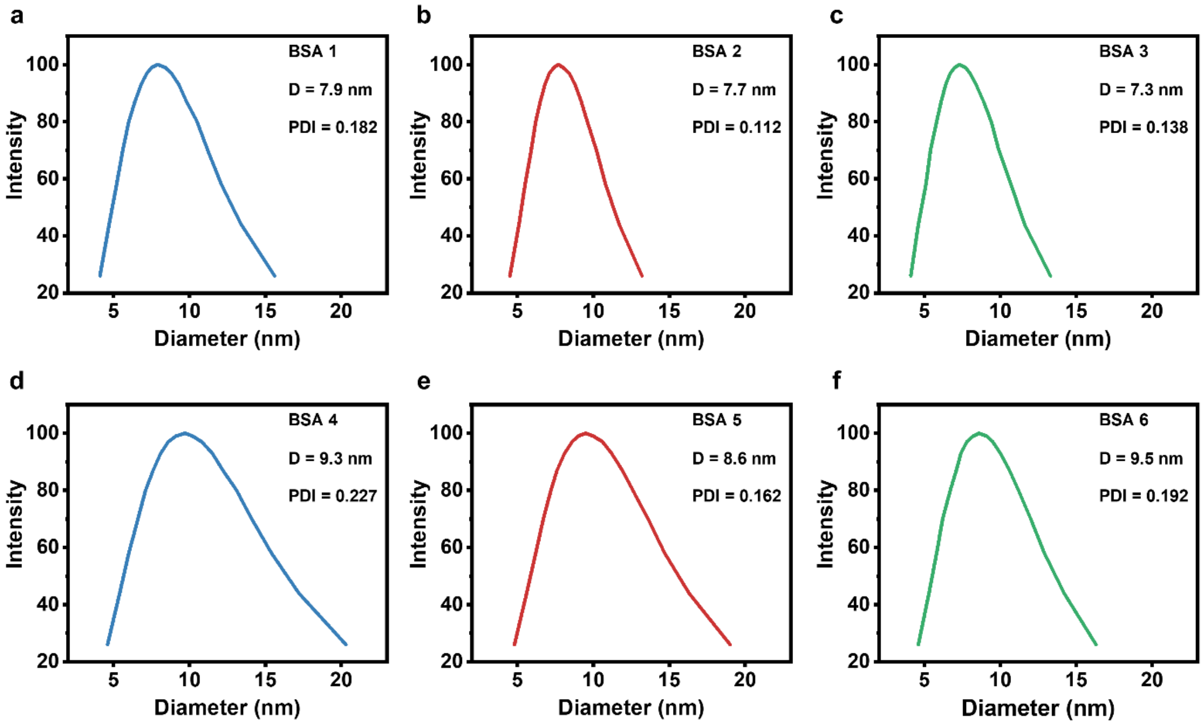


**Supplementary Figure 1. DLS characterization of BSA protein size distribution.**

**a-f,** DLS measurements of BSA proteins 1-6 at 25 °C (*n*=5 technical replicates). The mean hydrodynamic diameter (D) and polydispersity index (PDI) are indicated in each panel.

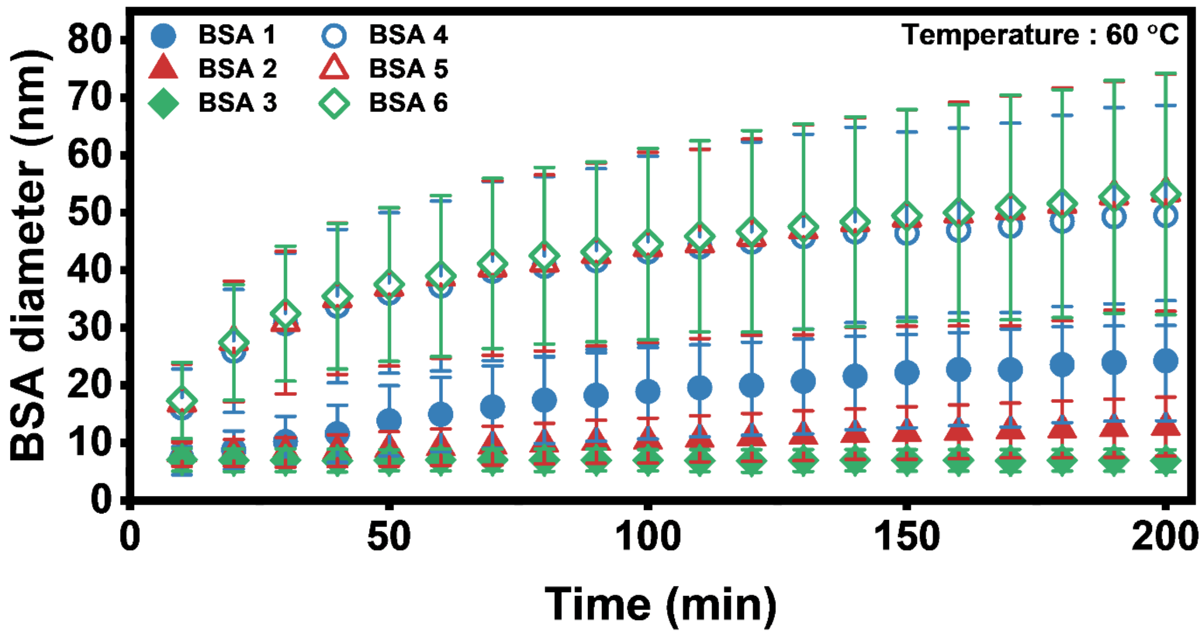


**Supplementary Figure 2. Time-dependent changes in protein size at 60 °C incubation temperature.**

Time-dependent changes in the mean hydrodynamic diameters of BSA proteins 1-6 at 60 °C. Values are presented as mean ± s.d. (*n*=5 technical replicates) where s.d. is defined as full-width-at-half-maximum (FWHM)/2.355.

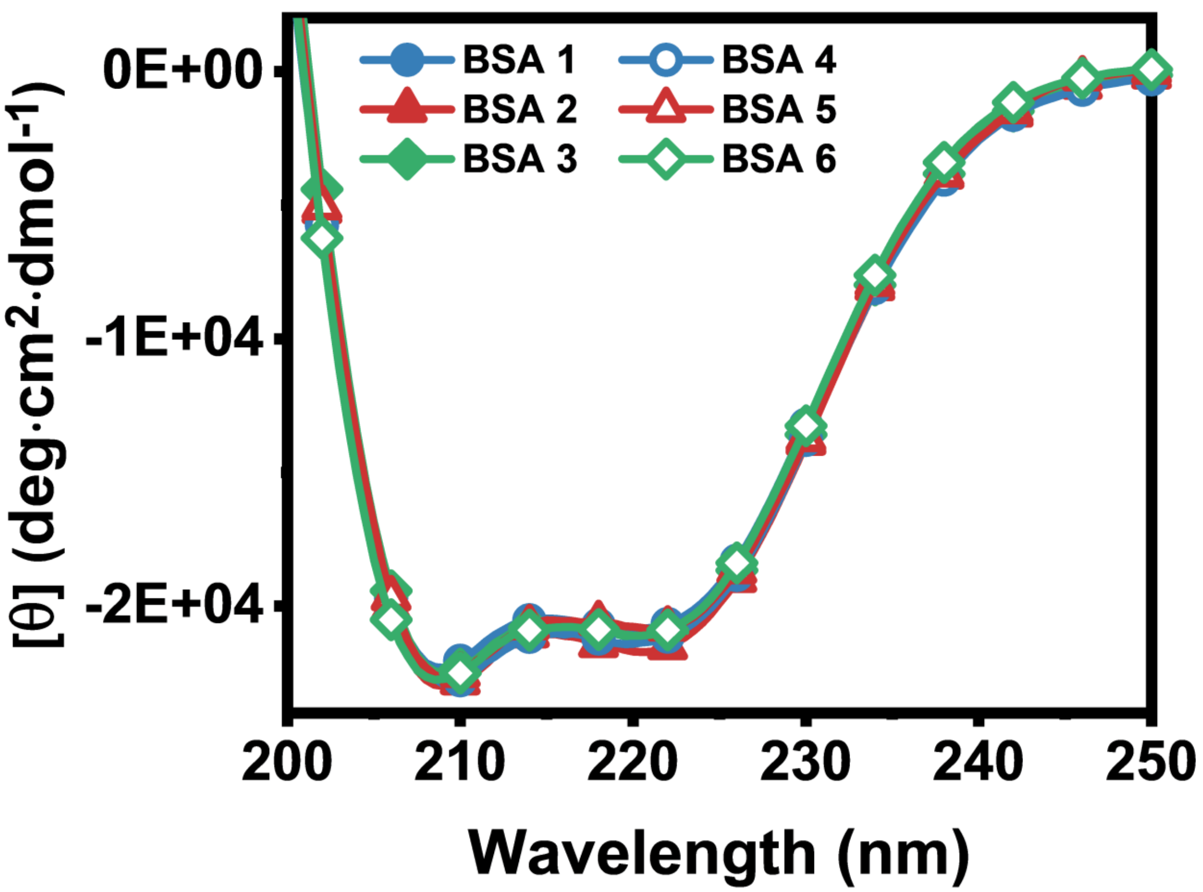


**Supplementary Figure 3. CD spectroscopy characterization of BSA proteins.**

Circular dichroism (CD) spectra of BSA proteins 1-6 at 25 °C are reported in molar residue ellipticity units [θ] (*n*=3 technical replicates).

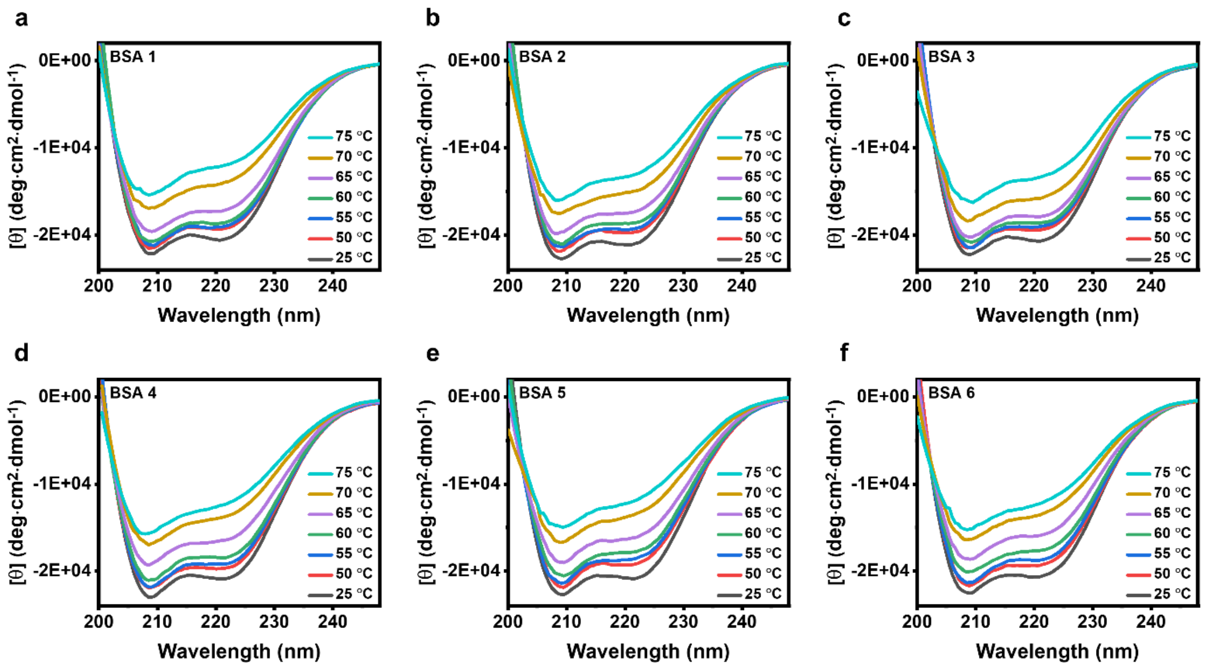


**Supplementary Figure 4. Effect of temperature on BSA secondary structure as determined by CD spectroscopy.**

**a-f,** CD spectra of BSA proteins 1-6 at 25, 50, 55, 60, 65, 70 and 75 °C are reported in molar residue ellipticity units [θ] (*n*=3 technical replicates). All elevated temperature measurements were recorded after a 5-min temperature equilibration period.

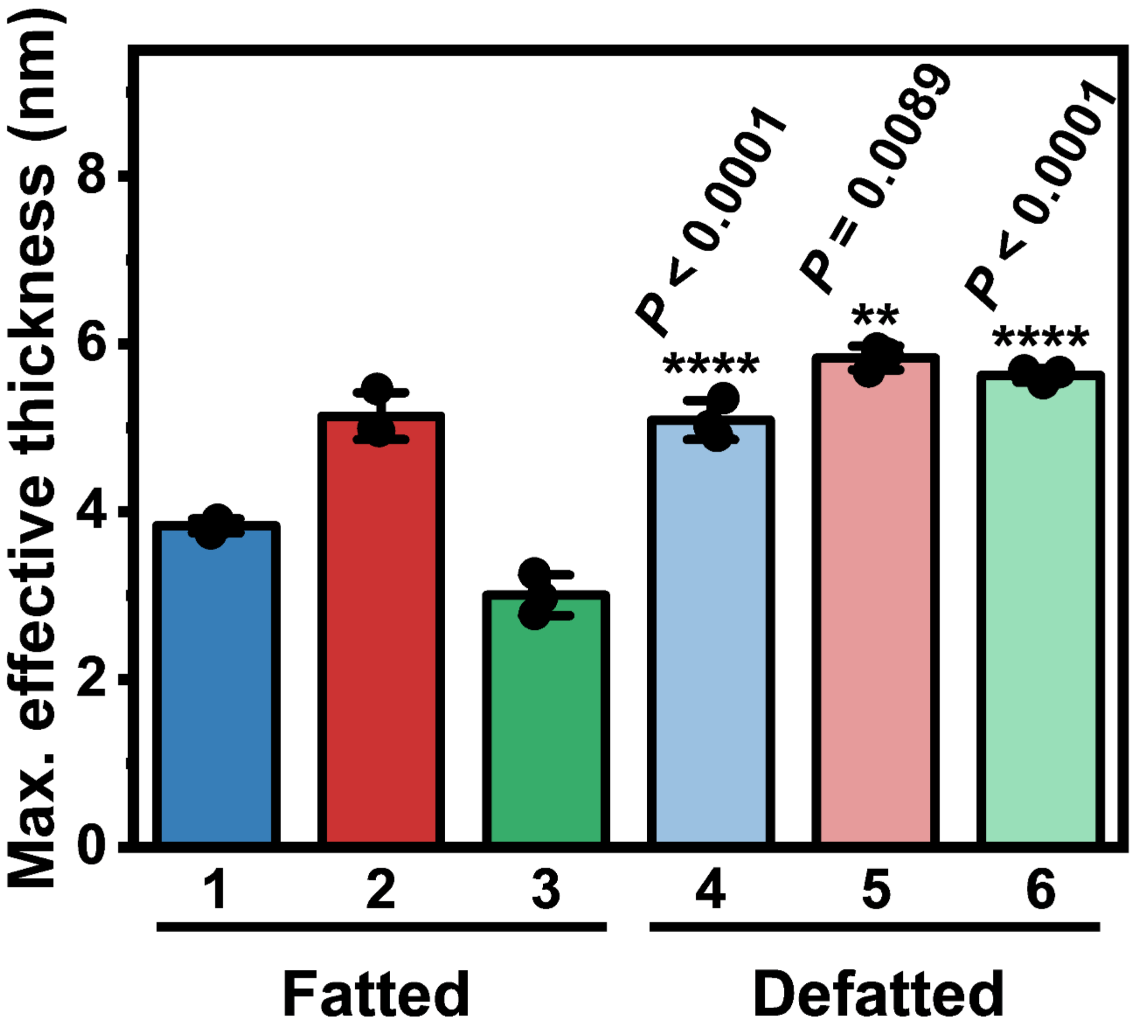


**Supplementary Figure 5. Effective thickness of adsorbed BSA protein layers based on viscoelastic fitting of QCM-D measurement data.**

The effective thickness of adsorbed layers comprising BSA proteins 1-6 was estimated based on Voigt-Voinova viscoelastic modeling of the QCM-D measurement data. The data are reported as mean ± s.d. (*n*=3 biological replicates, one-way analysis of variance (ANOVA) with Tukey's multiple comparisons test). *P* values are reported for defatted BSA samples (versus fatted BSA samples from the same fractionation method). Dots represent individual data points.

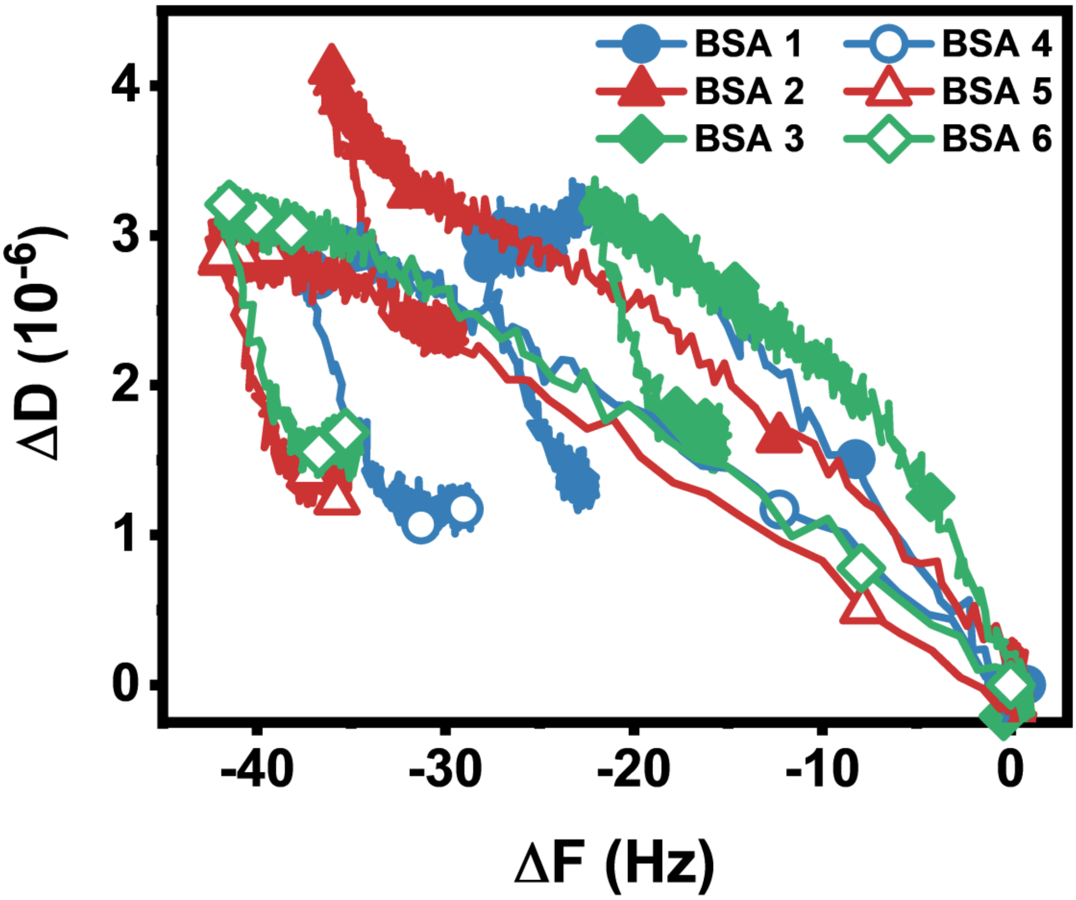


**Supplementary Figure 6. Comparison of QCM-D frequency-energy dissipation curves for BSA adsorption onto silica surfaces.**

Time-independent frequency-energy dissipation (F-D) curves derived from QCM-D frequency and energy dissipation shifts related to the adsorption of BSA proteins 1-6 onto silica surfaces at 25°C.

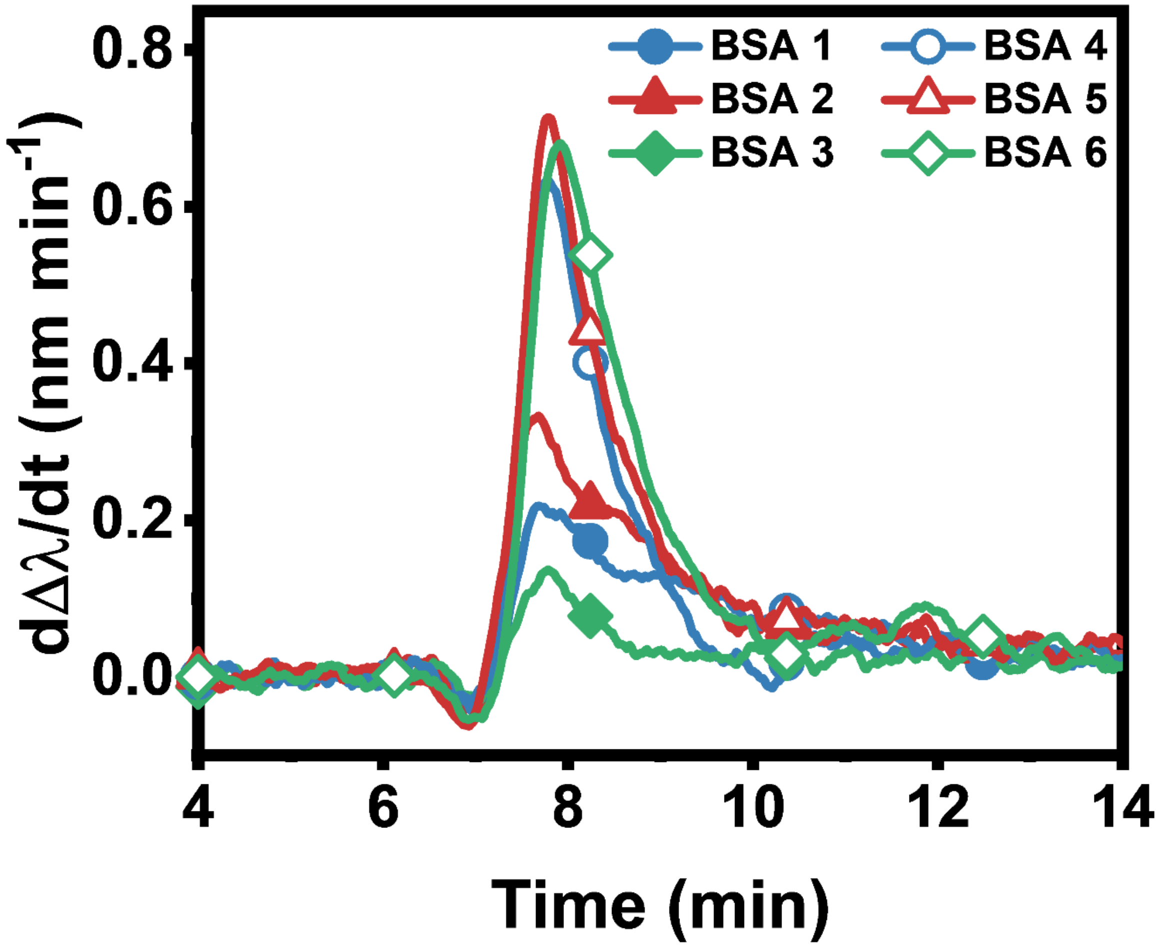


**Supplementary Figure 7. Time-derivative plot of LSPR-tracked BSA adsorption kinetics.**

Time-derivative of wavelength shifts (dΔλ/dt) corresponding to the LSPR-tracked adsorption of BSA proteins 1-6 onto silica-coated sensor surfaces.

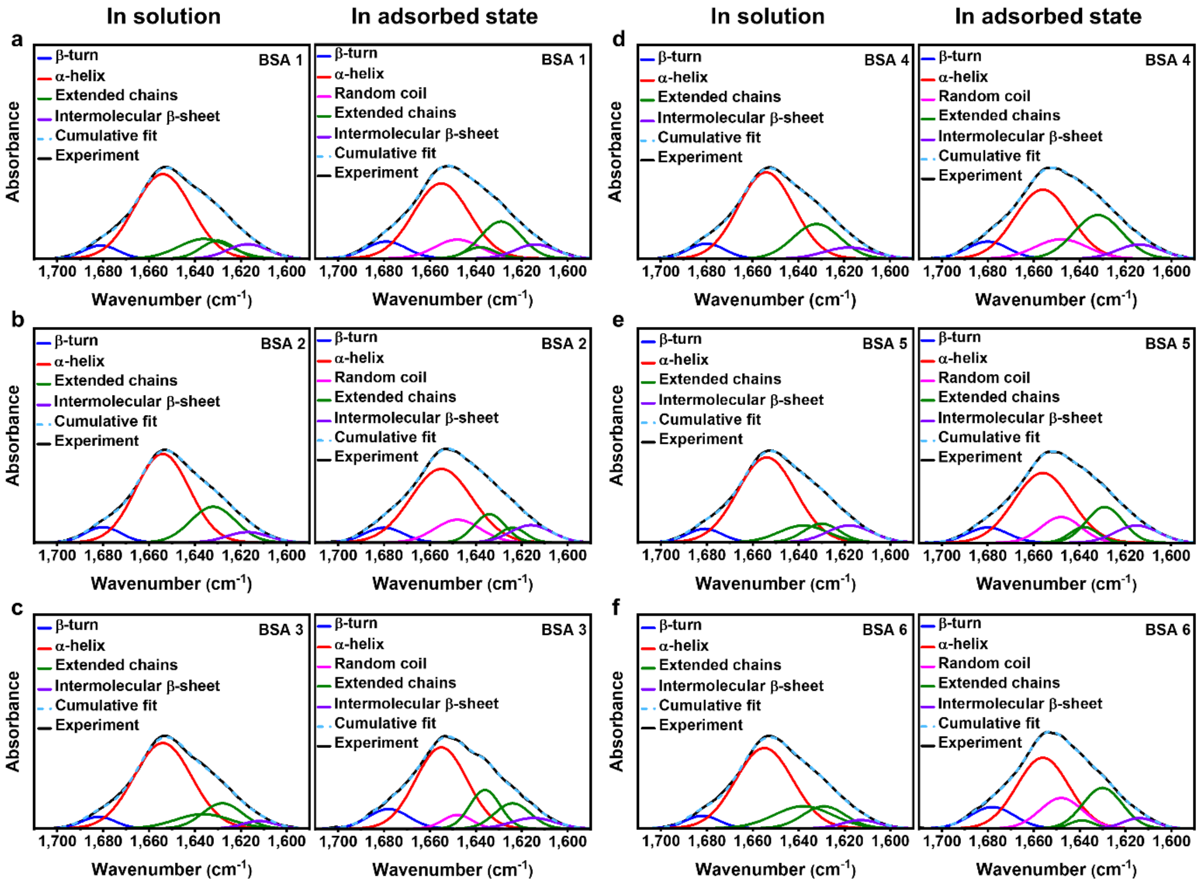


**Supplementary Figure 8. Quantification of adsorption-related protein conformational changes in the secondary structure of BSA proteins based on ATR-FTIR measurements.**

Amide I regions from attenuated total reflection-Fourier transform infrared (ATR-FTIR) spectrograms of (**a-f**) BSA proteins 1-6 in solution (left panels) and in the adsorbed state (right panels). Experimentally obtained absorbance spectra were resolved into individual component curves, which were each assigned to a secondary structure element.

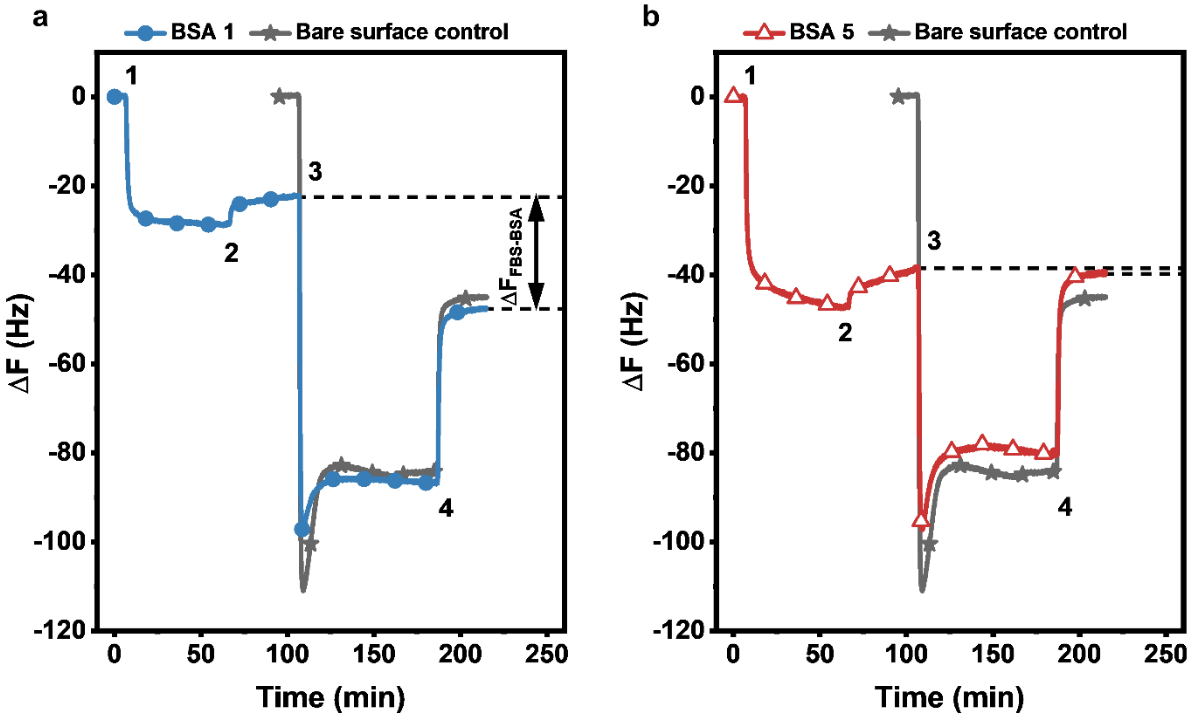


**Supplementary Figure 9. Quantification of BSA surface passivation performance against serum biofouling based on QCM-D measurements.**

Time-resolved QCM-D frequency shifts (ΔF) evaluating the blocking efficiency of BSA protein coatings against fetal bovine serum (FBS) fouling of silica surfaces for **a,** BSA protein 1 and **b,** BSA protein 5. The protocol steps involved (1) 100 µM BSA addition, (2) buffer washing step, (3) addition of undiluted FBS, and (4) a buffer washing step. Bare surface control refers to a control experiment whereby FBS was added to an uncoated silica surface (no BSA coating). The difference in ΔF values due to BSA adsorption alone (post-washing) and after FBS incubation (post-washing) was computed ($\Delta F_{FBS\text{-}BSA}$, indicated by dashed lines) and the blocking efficiency percentage was calculated from this value compared to the equivalent value computed in the control experiment without BSA coating (see Methods section for more information).

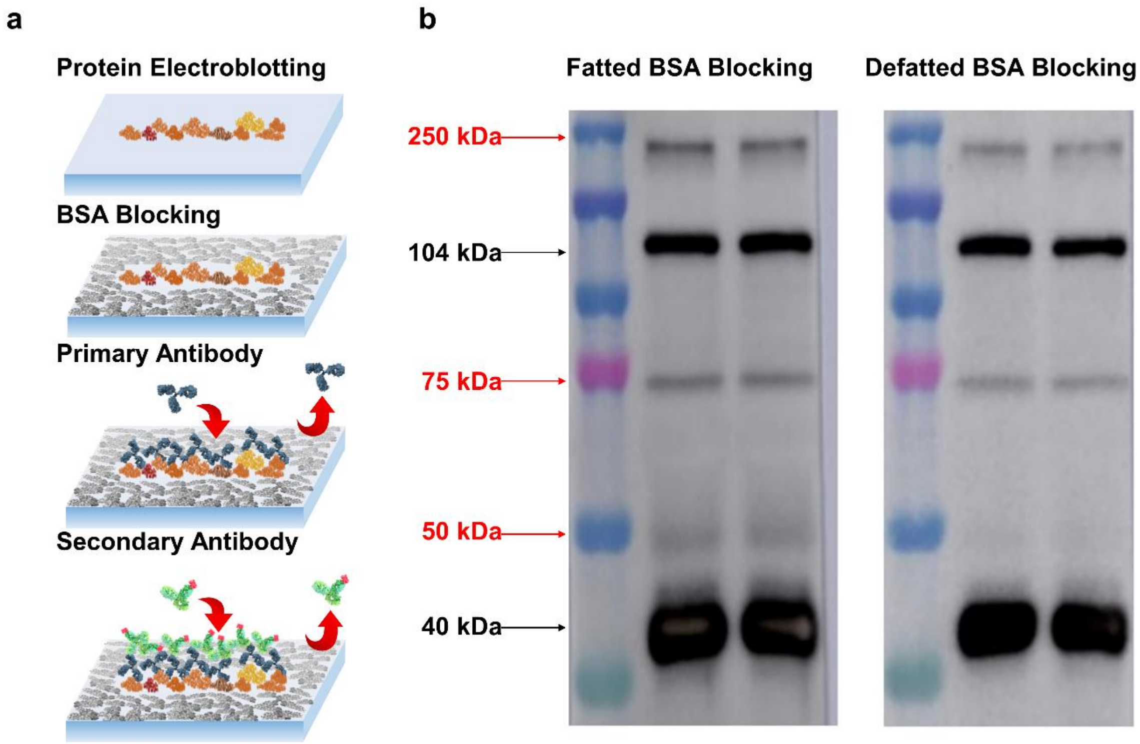


**Supplementary Figure 10. Blocking performance of BSA protein reagents in Western blot experiments.**

**a,** Schematic illustration of the BSA blocking step in a Western blot experiment. **b,** Three Western blot lanes consisted of a colored molecular weight marker (lane 1) and normal human serum (NHS) samples (lanes 2-3). Each membrane was individually blocked with BSA 1 (fatted) or BSA 5 (defatted) prior to C3b antibody incubation. Red arrows indicate the position of nonspecific bands located near the 250, 75, and 50 kDa molecular weight markers. Black arrows indicate the position of specific bands corresponding to C3b (104 kDa) and iC3b (40 kDa). The objective of the blocking step is to minimize signal noise coming from nonspecific bands in order to focus analysis on specific bands.

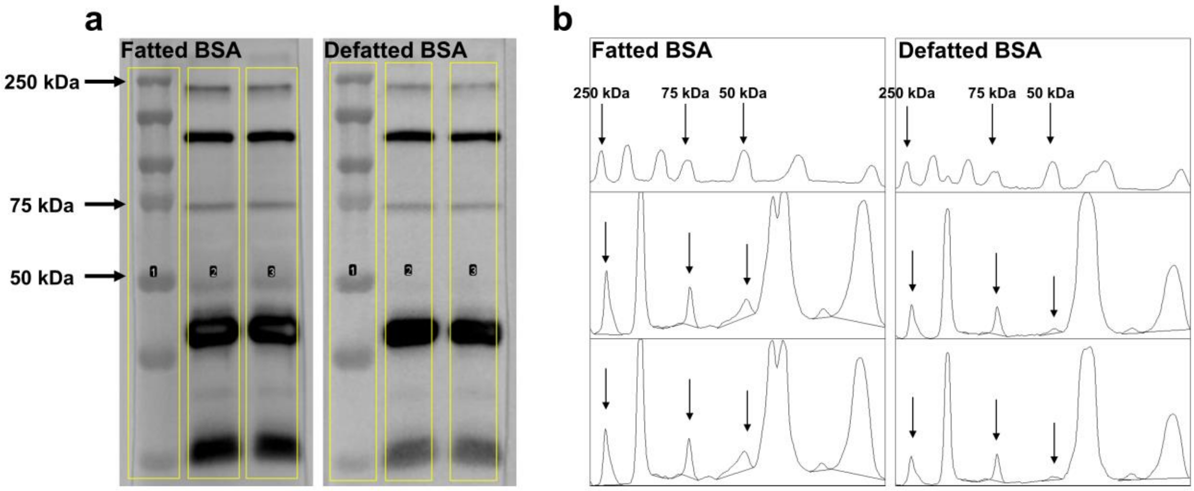


**Supplementary Figure 11. Quantification of Western blot band intensities.**

**a,** Greyscale digital images of Western blots with the three lanes selected for the extraction of lane intensity profiles. Lane 1 is from molecular weight markers while lanes 2 and 3 are from NHS samples. Left and right panels show membranes blocked by BSA 1 and BSA 5 proteins, respectively. **b,** Intensity profiles from lanes 1, 2, and 3 are shown in the top, middle, and bottom rows, respectively. Bands in the digital image appear as peaks in the intensity profile. Baselines were established by connecting the two minima on both sides of a selected peak. The intensity values were obtained based on the area under the curves bounded by the established baseline. Left and right panels show membranes blocked by BSA proteins 1 and 5, respectively. Arrows indicate the position of nonspecific bands located near the 250, 75, and 50 kDa molecular weight markers.

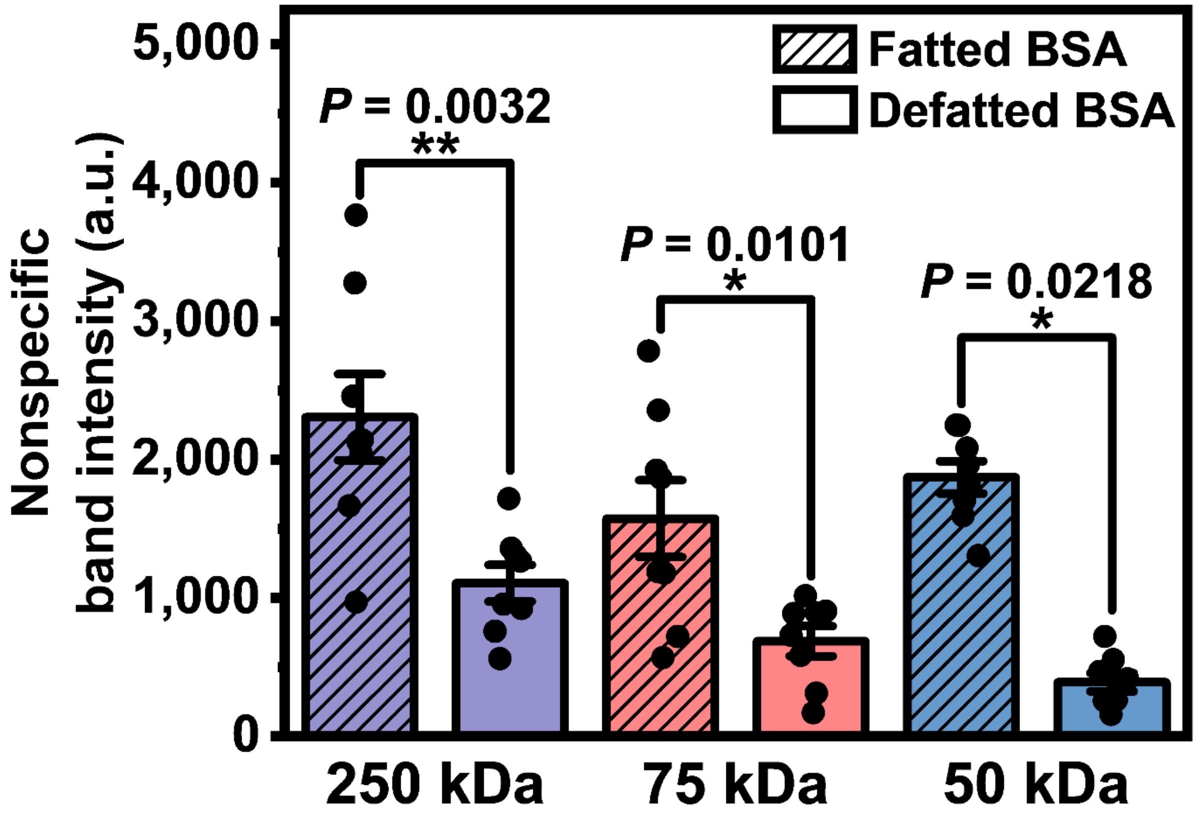


**Supplementary Figure 12. Comparison of nonspecific band intensities resulting from the use of fatted and defatted BSA proteins in Western blot experiments.**

Intensity values of the nonspecific bands that are located near the 250, 75, and 50 kDa molecular weight markers from membranes that were blocked with a blocking solution that included BSA 1 (fatted) or BSA 5 (defatted) (*n*=8 biological replicates, unpaired t-test). Data were obtained from Fiji/ImageJ software program and are reported as mean ± s.e.m. Dots represent individual data points and intensities are in arbitrary units.

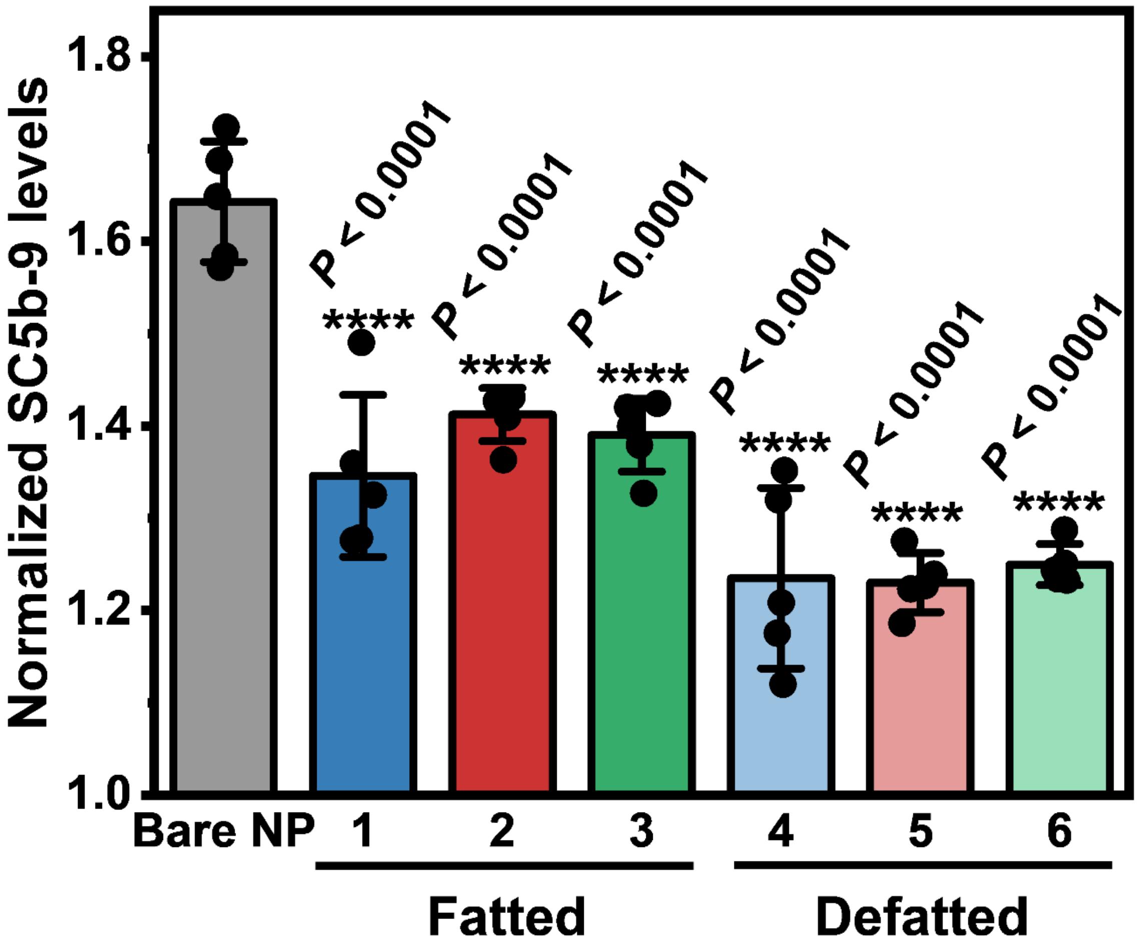


**Supplementary Figure 13. Reduction in silica nanoparticle-induced complement activation due to BSA coatings, as quantified by normalized SC5b-9 levels.**

The level of complement activation for each sample was determined by measuring SC5b-9 concentrations by enzyme-linked immunosorbent assay (ELISA) and normalized against SC5b-9 levels in NHS without nanoparticles [*n*=5 biological replicates, one-way ANOVA with Dunnett's multiple comparisons test (versus uncoated nanoparticles)]. Data are reported as mean ± s.e.m. and dots represent individual data points.

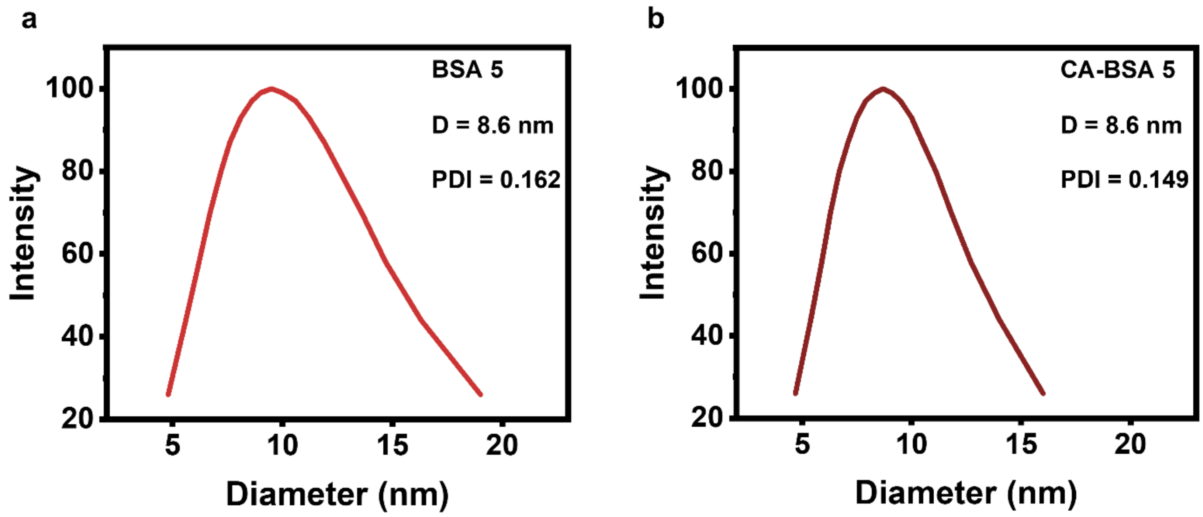


**Supplementary Figure 14. DLS characterization of BSA 5 size distribution without and with caprylic acid doping.**

DLS measurements of **a,** BSA 5 and **b,** caprylic acid-doped BSA 5 (CA-BSA 5) at 25 °C (*n*=5 technical replicates). The mean hydrodynamic diameter (D) and polydispersity index (PDI) are indicated in each panel.

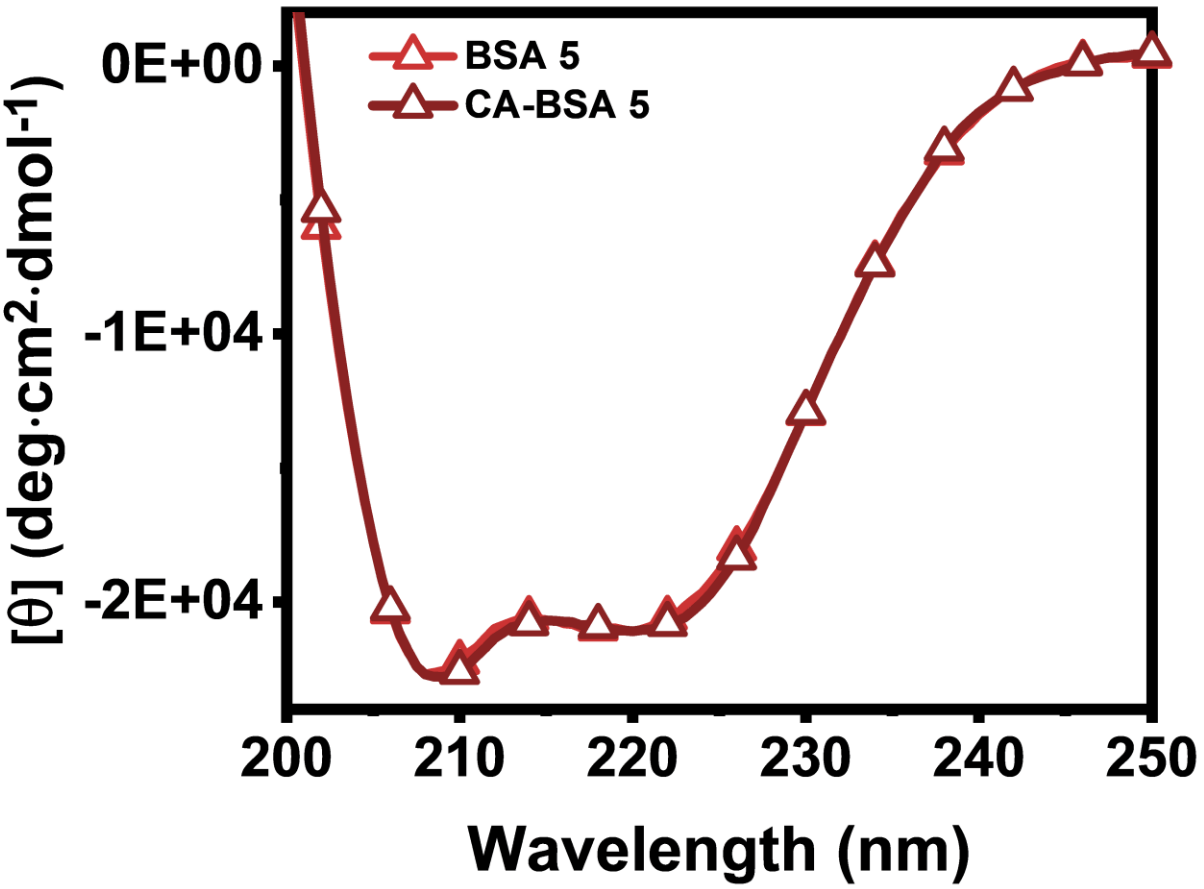


**Supplementary Figure 15. CD spectroscopy characterization of BSA 5 without and with caprylic acid doping.**

Circular dichroism (CD) spectra of BSA 5 and caprylic acid-doped BSA 5 (CA-BSA 5) at 25 °C are reported in molar residue ellipticity units [θ] (*n*=3 technical replicates).

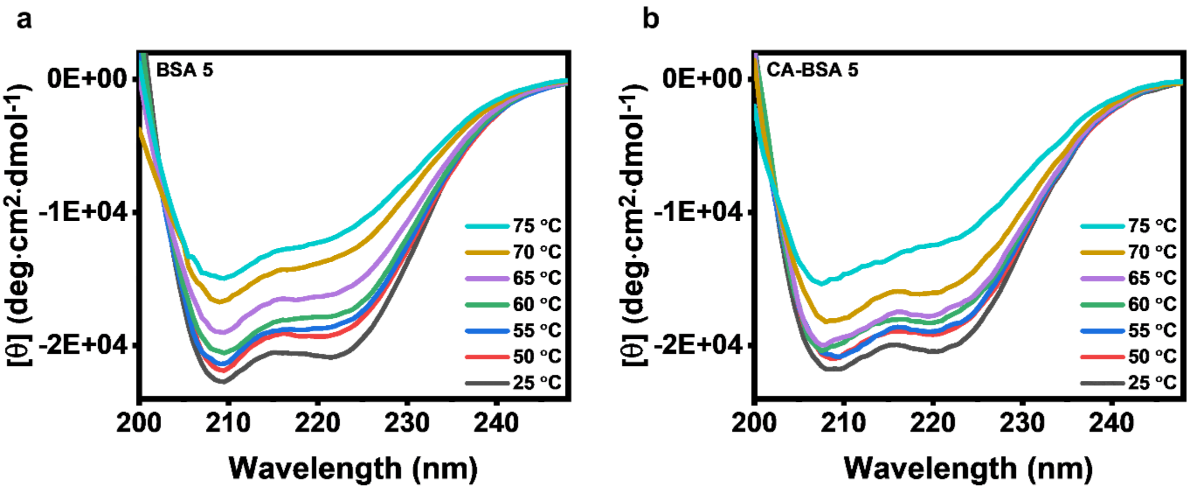


**Supplementary Figure 16. Effect of temperature on secondary structure of BSA 5 without and with caprylic acid doping as determined by CD spectroscopy.**

CD spectra of **a,** BSA 5 and **b,** caprylic acid-doped BSA 5 (CA-BSA 5) at 25, 50, 55, 60, 65, 70 and 75 °C are reported in molar residue ellipticity units [θ] ($n$=3 technical replicates). All elevated temperature measurements were recorded after a 5-min temperature equilibration period.

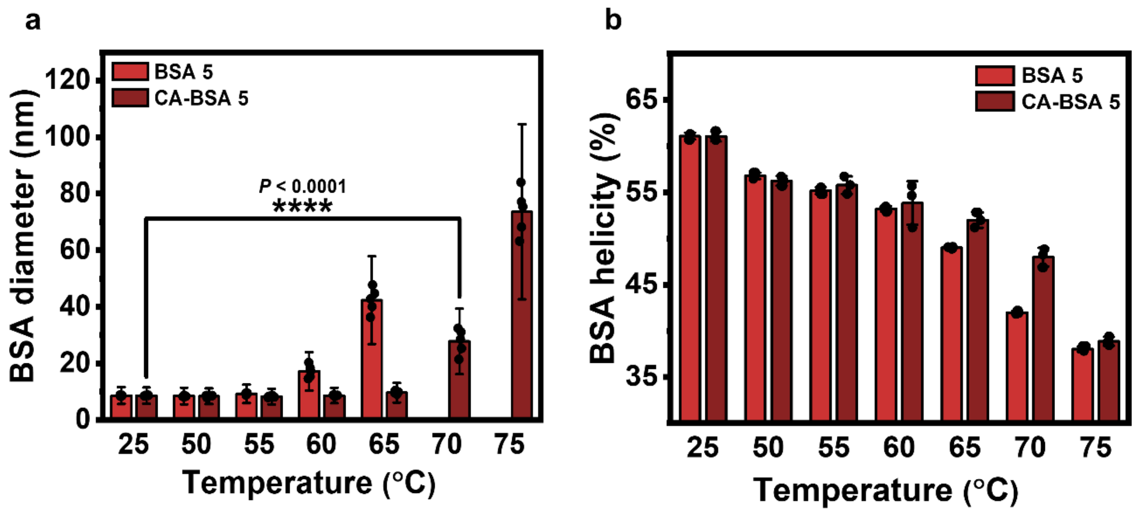


**Supplementary Figure 17. Temperature profiling of BSA conformational stability in solution for BSA 5 without and with caprylic acid doping.**

**a,** Hydrodynamic diameter of BSA 5 and caprylic acid-doped BSA 5 (CA-BSA 5) as a function of temperature, as measured in DLS experiments. Data are reported as mean ± s.d. [$n$=5 technical replicates, one-way ANOVA with Dunnett's multiple comparisons test (versus data at 25 °C) for CA-BSA 5 data]. Dots represent individual data points. **b,** Fractional percentage of α-helicity in BSA 5 and CA-BSA 5 protein molecules as a function of temperature, as measured in CD spectroscopy experiments. Mean values are presented on top of each column. Values were computed from molar residue ellipticity data and data are reported as mean ± s.d. ($n$=3 technical replicates). Dots represent individual data points.

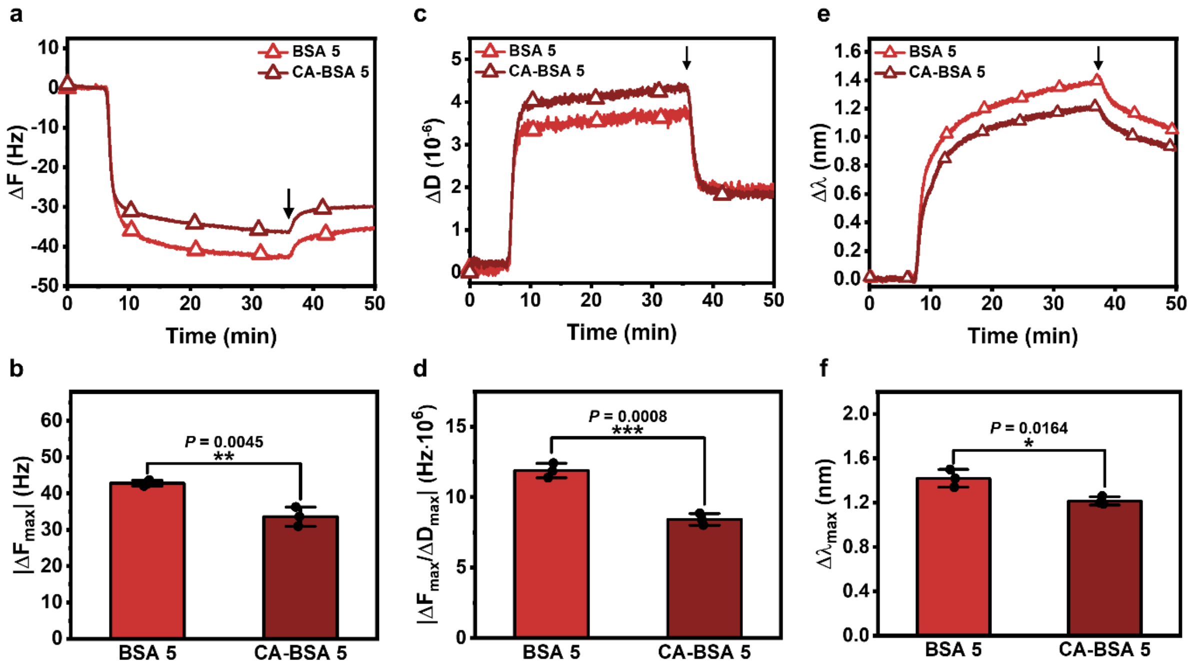


**Supplementary Figure 18. Real-time monitoring of BSA protein adsorption onto silica surfaces for BSA 5 without and with caprylic acid doping.**

**a,** Time-resolved QCM-D ΔF shifts and **b,** corresponding $|\Delta F_{max}|$ shifts at saturation. **c,** Time-resolved QCM-D ΔD shifts. **d,** $|\Delta F_{max}/\Delta D_{max}|$ ratios obtained from saturation data in panels **a** and **c**. **e-f,** LSPR experiments were conducted to measure $\Delta\lambda_{max}$ signals related to the protein adsorption process. **e,** Time-resolved LSPR wavelength shifts (Δλ) and **f,** corresponding $\Delta\lambda_{max}$ shifts at saturation. Data in **b, d, f** are reported as mean ± s.d. (*n*=3 biological replicates, unpaired t-test). Dots represent individual data points.

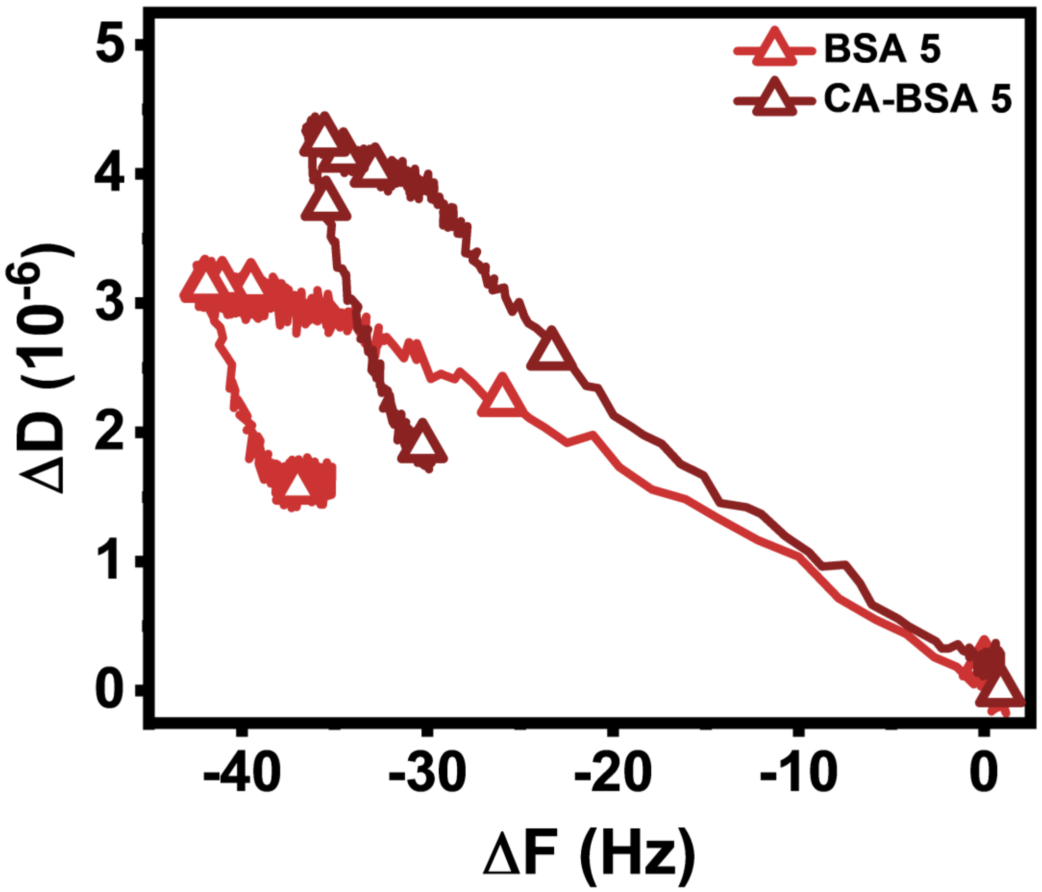


**Supplementary Figure 19. Comparison of QCM-D frequency-energy dissipation curves for comparing the adsorption of BSA 5 without and with caprylic acid doping onto silica surfaces.**

Time-independent frequency-energy dissipation (F-D) curves derived from QCM-D frequency and energy dissipation shifts related to the adsorption of BSA 5 and caprylic acid-doped BSA 5 (CA-BSA 5) onto silica surfaces at 25°C.

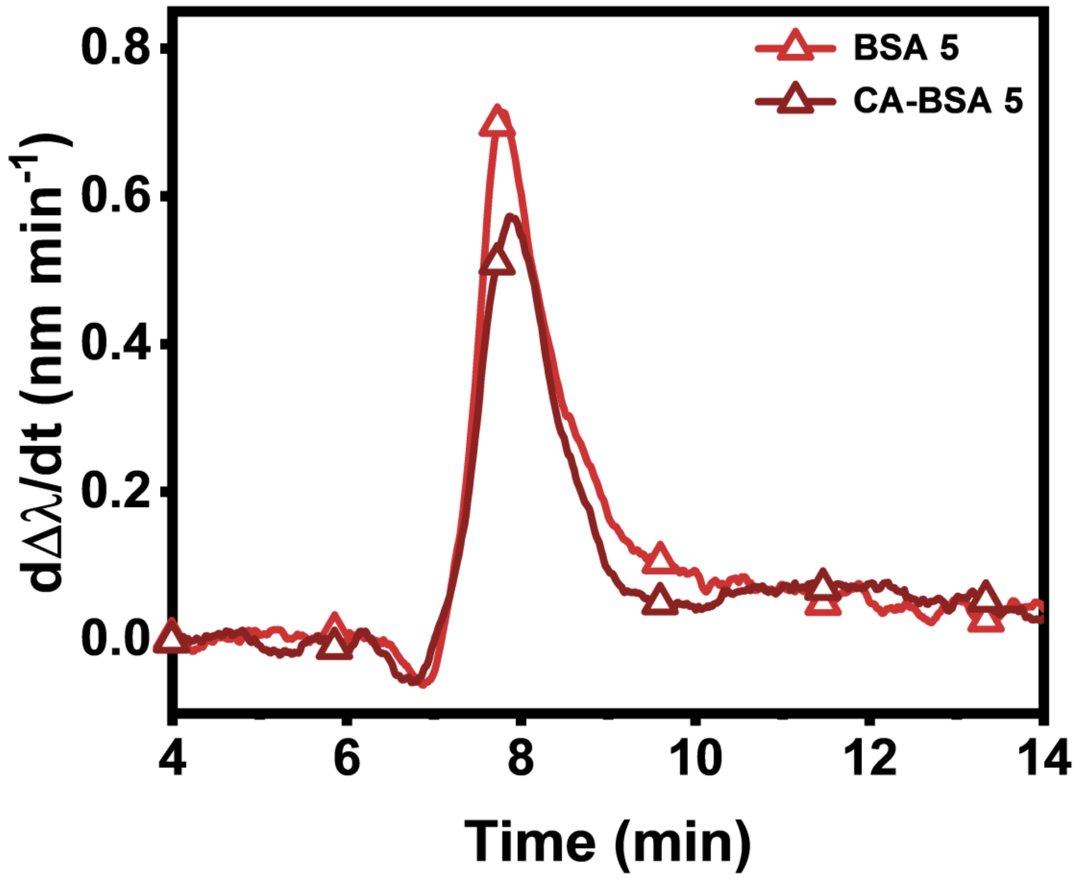


**Supplementary Figure 20. Time-derivative plot of LSPR-tracked BSA adsorption kinetics for BSA 5 without and with caprylic acid doping.**

Time-derivative of wavelength shifts (dΔλ/dt) corresponding to the LSPR-tracked adsorption of BSA 5 and caprylic acid-doped BSA 5 (CA-BSA 5) onto silica-coated sensor surfaces.

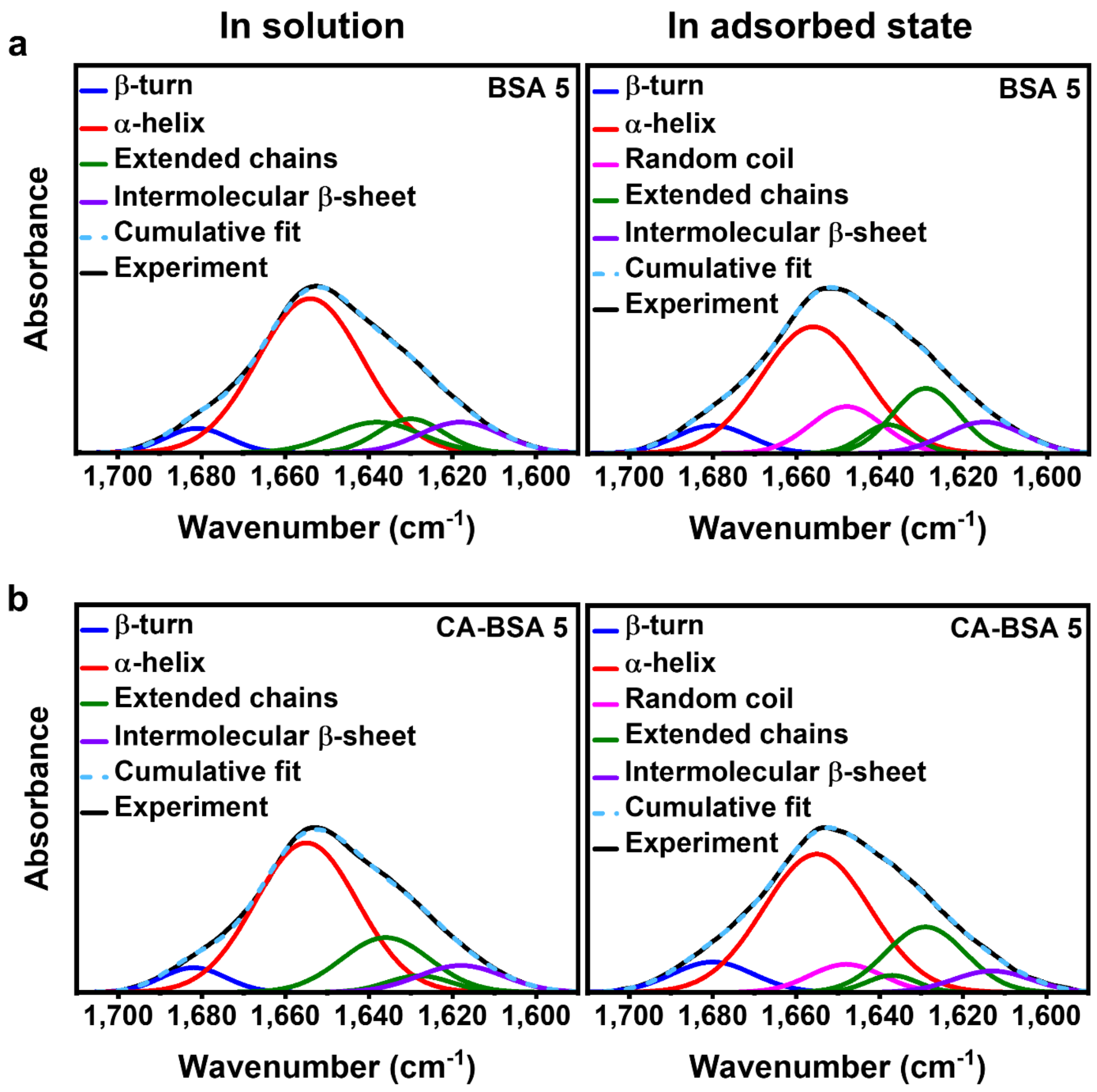


**Supplementary Figure 21. Comparison of adsorption-related protein conformational changes in the secondary structure of BSA 5 without and with caprylic acid doping based on ATR-FTIR measurements.**

Amide I regions from attenuated total reflection-Fourier transform infrared (ATR-FTIR) spectrograms of **a,** BSA 5 and **b,** caprylic acid-doped BSA 5 (CA-BSA 5) in solution (left panels) and in the adsorbed state (right panels). Experimentally obtained absorbance spectra were resolved into individual component curves, which were each assigned to a secondary structure element.

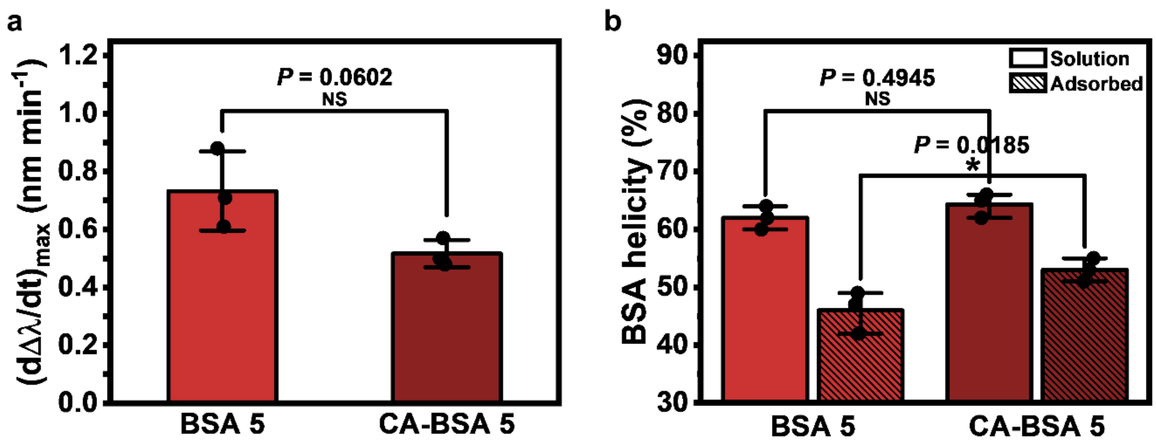


**Supplementary Figure 22. Quantitative evaluation of adsorption-related conformational changes in BSA protein structure for BSA 5 without and with caprylic acid doping.**

**a,** Maximum rate of change in the LSPR wavelength shift $(d\Delta\lambda/dt)_{max}$ during the initial adsorption stage of BSA 5 and caprylic acid-doped BSA 5 (CA-BSA 5). Values are computed from data in Supplementary Figure 18. Data are reported as mean ± s.d. ($n$=3 biological replicates, unpaired t-test). **b,** Fractional percentage of α-helicity in BSA 5 and CA-BSA 5 protein molecules in solution and in the adsorbed state, as determined in ATR-FTIR spectroscopy experiments. Data are reported as mean ± s.d. ($n$=3 biological replicates, two-way ANOVA with Sidak's multiple comparisons test). Dots represent individual data points.

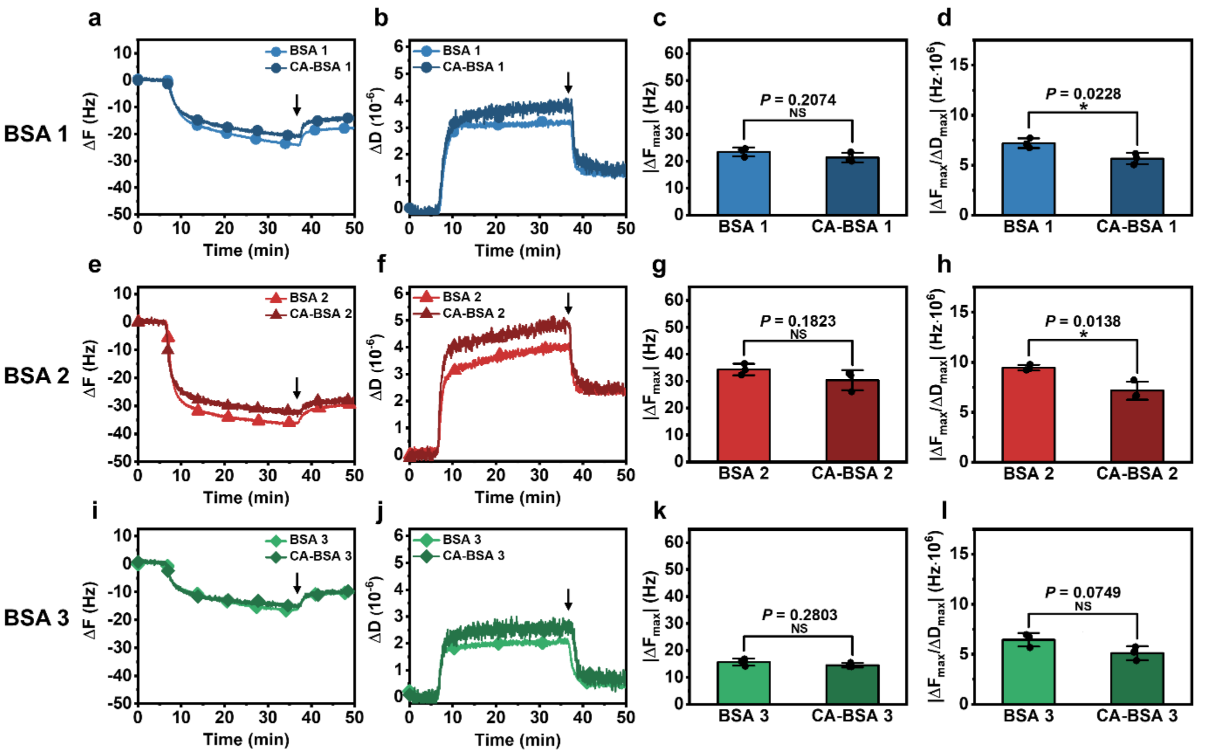


**Supplementary Figure 23. Influence of caprylic acid doping on the adsorption of fatted BSA proteins 1-3, as measured by QCM-D experiments.**

**(a-f) a,** Time-resolved QCM-D ΔF and **b,** ΔD shifts along with **c,** corresponding $|\Delta F_{max}|$ shifts at saturation (from panel **a**) and **d,** $|\Delta F_{max}/\Delta D_{max}|$ ratios obtained from saturation data in panels **a** and **b** for BSA 1 without and with caprylic acid doping (labeled as BSA 1 and CA-BSA 1, respectively). **(e-h)** Equivalent data for BSA 2. **(i-l)** Equivalent data for BSA 3. Data in **c, d, g, h, k, l** are reported as mean ± s.d. (*n*=3 biological replicates, unpaired t-test). Dots represent individual data points.

**Supplementary Table 1. Summary of BSA protein reagents.**

The albumin content percentage by agarose electrophoresis and the percentage of free fatty acids were obtained from the manufacturer's certificates of analysis. The amount of free fatty acid in BSA 1-3 were not reported because they were "fatted" reagents while the free fatty acid percentage for BSA 4-6 verifies that they were "defatted" reagents.

| BSA name | Catalogue no. | Lot no. | Purification method | Albumin by agarose electrophoresis (%) | Free fatty acid (%) |
|---|---|---|---|---|---|
| BSA 1 | A2153 | SLBW1397 | Cold ethanol fractionation | 100 | NA |
| BSA 2 | A3059 | SLBS4332 | Heat shock fractionation | 100 | NA |
| BSA 3 | A7638 | SLBW3326 | Cold ethanol and heat shock fractionation | 100 | NA |
| BSA 4 | A6003 | SLBR4436V | Cold ethanol fractionation | 100 | 0.00 |
| BSA 5 | A7030 | SLBK3074V | Heat shock fractionation | 99 | 0.00 |
| BSA 6 | A0281 | 021M7403V | Cold ethanol and heat shock fractionation | 100 | 0.01 |

**Supplementary Table 2. Helicity percentage of BSA proteins 1-6 in solution at 25, 50, 55, 60, 65, 70, and 75 °C from CD spectroscopy measurements.**

Data are reported as mean ± s.d. (*n*=3 technical replicates).

| **BSA Type** | **25 °C (%)** | **50 °C (%)** | **55 °C (%)** | **60 °C (%)** | **65 °C (%)** | **70 °C (%)** | **75 °C (%)** |
|---|---|---|---|---|---|---|---|
| BSA 1 | 60.7 ± 0.3 | 57.0 ± 0.3 | 56.2 ± 0.1 | 55.2 ± 0.3 | 51.3 ± 0.6 | 43.2 ± 0.3 | 38.5 ± 0.3 |
| BSA 2 | 62.9 ± 0.2 | 58.0 ± 0.3 | 57.0 ± 0.4 | 55.3 ± 0.1 | 51.9 ± 0.3 | 46.0 ± 0.8 | 41.0 ± 0.2 |
| BSA 3 | 61.5 ± 0.5 | 57.1 ± 0.4 | 56.2 ± 0.3 | 55.2 ± 0.2 | 53.3 ± 0.6 | 47.6 ± 0.5 | 41.5 ± 0.5 |
| BSA 4 | 62.0 ± 0.2 | 58.1 ± 0.2 | 56.7 ± 0.3 | 54.8 ± 0.2 | 49.8 ± 0.3 | 42.8 ± 0.5 | 39.0 ± 0.4 |
| BSA 5 | 61.1 ± 0.3 | 56.8 ± 0.2 | 55.1 ± 0.4 | 53.2 ± 0.3 | 48.9 ± 0.1 | 42.1 ± 0.1 | 38.0 ± 0.2 |
| BSA 6 | 61.3 ± 0.5 | 56.8 ± 0.2 | 55.4 ± 0.2 | 52.6 ± 0.3 | 48.1 ± 0.5 | 41.7 ± 0.5 | 38.3 ± 0.1 |

**Supplementary Table 3. Helicity percentage of BSA proteins 1-6 in solution and in the adsorbed state from ATR-FTIR spectroscopy measurements.**

Data are reported as mean ± s.d. (*n*=3 biological replicates).

| **BSA type** | **Solution (%)** | **Adsorbed (%)** |
|---|---|---|
| BSA 1 | 63.7 ± 1.5 | 52.7 ± 3.1 |
| BSA 2 | 65.3 ± 0.6 | 53.7 ± 2.1 |
| BSA 3 | 65.3 ± 1.5 | 54 ± 2.7 |
| BSA 4 | 61.7 ± 1.2 | 45.7 ± 3.5 |
| BSA 5 | 62.0 ± 2.0 | 46.0 ± 3.6 |
| BSA 6 | 59.7 ± 1.5 | 44.7 ± 3.2 |

**Supplementary Table 4. Helicity percentage of BSA 5 and CA-BSA 5 in solution at 25, 50, 55, 60, 65, 70, and 75 °C from CD spectroscopy measurements.**

Data are reported as mean ± s.d. (*n*=3 technical replicates).

| **BSA type** | **25 °C (%)** | **50 °C (%)** | **55 °C (%)** | **60 °C (%)** | **65 °C (%)** | **70 °C (%)** | **75 °C (%)** |
|---|---|---|---|---|---|---|---|
| BSA 5 | 61.1 ± 0.4 | 56.8 ± 0.3 | 55.2 ± 0.4 | 53.2 ± 0.3 | 49.0 ± 0.1 | 42.0 ± 0.2 | 38.0 ± 0.3 |
| CA-BSA 5 | 61.0 ± 0.5 | 56.2 ± 0.5 | 55.3 ± 0.5 | 53.5 ± 2.0 | 52.0 ± 0.8 | 48.0 ± 1.0 | 38.9 ± 0.5 |

**Supplementary Table 5. Helicity percentage of BSA 5 and CA-BSA 5 in solution and in the adsorbed state from ATR-FTIR spectroscopy measurements.**

Data are reported as mean ± s.d. (*n*=3 biological replicates).

| BSA type | Solution (%) | Adsorbed (%) |
|---|---|---|
| BSA 5 | 62.0 ± 2.0 | 46.0 ± 3.6 |
| CA-BSA 5 | 64.3 ± 2.1 | 53.0 ± 2.0 |

**Supplementary Table 6. Details of statistical analyses.**

The type of multiple comparisons test conducted after ANOVA are listed in the “Multiple comparisons test” column. F-values from one-way or two-way ANOVA and t-values from unpaired t-tests are listed in the “F, t values” column. Multiplicity-adjusted *P* values from multiple comparisons tests and *P* values from unpaired t-tests are listed in the “*P* value” column.

| **Figure** | **Analysis** | **Multiple comparisons test** | **F, t values** | ***P* value** |
|---|---|---|---|---|
| Fig. 2a, BSA 1 | One-way ANOVA | Dunnett's test, control: 25 °C | F (4, 20) = 20.46 | *P*<0.0001 |
| Fig. 2a, BSA 2 | One-way ANOVA | Dunnett's test, control: 25 °C | F (5, 24) = 14.42 | *P*<0.0001 |
| Fig. 2a, BSA 3 | One-way ANOVA | Dunnett's test, control: 25 °C | F (6, 28) = 13.48 | *P*<0.0001 |
| Fig. 2a, BSA 4 | One-way ANOVA | Dunnett's test, control: 25 °C | F (4, 20) = 13.57 | *P*<0.0001 |
| Fig. 2a, BSA 5 | One-way ANOVA | Dunnett's test, control: 25 °C | F (4, 20) = 16.78 | *P*<0.0001 |
| Fig. 2a, BSA 6 | One-way ANOVA | Dunnett's test, control: 25 °C | F (4, 20) = 16.32 | *P*<0.0001 |
| Fig. 2a, BSA 1 – 6 at 25 °C | One-way ANOVA | Tukey's test | F (5, 24) = 0.5274 | *P*=0.7532 |
| Fig. 3d | One-way ANOVA | Tukey's test | F (5, 12) = 126.0 | *P*<0.0001 |
| Fig. 3e | One-way ANOVA | Tukey's test | F (5, 12) = 37.27 | *P*<0.0001 |
| Fig. 3f | One-way ANOVA | Tukey's test | F (5, 12) = 34.01 | *P*<0.0001 |
| Fig. 4a | One-way ANOVA | Tukey's test | F (5, 12) = 38.36 | *P*<0.0001 |
| Fig. 4b | Two-way ANOVA | Tukey's test | BSA type: F (5, 24) = 11.33 | *P*<0.0001 |
| | | | Solution vs adsorbed: F (1, 24) = 285.3 | *P*<0.0001 |
| | | | Interaction: F (5, 24) = 1.516 | *P*=0.2221 |
| Fig. 5c | One-way ANOVA | Tukey's test | F (5, 12) = 18.83 | *P*<0.0001 |
| Fig. 5d | One-way ANOVA | Tukey's test | F (5, 24) = 12.19 | *P*<0.0001 |
| Supplementary Fig. 6 | One-way ANOVA | Tukey's test | F (5, 12) = 97.55 | *P*<0.0001 |
| Supplementary Fig. 12, 250 kDa | Unpaired t-test | | t=3.551, df=14 | *P*=0.0032 |
| Supplementary Fig. 12, 75 kDa | Unpaired t-test | | t=2.972, df=14 | *P*=0.0101 |

| | | | | |
|---|---|---|---|---|
| Supplementary Fig. 12, 50 kDa | Unpaired t-test | | t=2.842, df=8 | *P*=0.0218 |
| Supplementary Fig. 13 | One-way ANOVA | Dunnett's test, control: Bare NP | F (6, 28) = 29.43 | *P*<0.0001 |
| Supplementary Fig. 17a, CA-BSA 5 | One-way ANOVA | Dunnett's test, control: 25 °C | F (5, 24) = 10.26 | *P*<0.0001 |
| Supplementary Fig. 17a, BSA 5 and CA-BSA 5 at 25 °C | Unpaired t-test | | t=0.01079, df=8 | *P*=0.9917 |
| Supplementary Fig. 18d | Unpaired t-test | | t=5.755, df=4 | *P*=0.0045 |
| Supplementary Fig. 18e | Unpaired t-test | | t=9.056, df=4 | *P*=0.0008 |
| Supplementary Fig. 18f | Unpaired t-test | | t=3.979, df=4 | *P*=0.0164 |
| Supplementary Fig. 22a | Unpaired t-test | | t=2.598, df=4 | *P*=0.0602 |
| Supplementary Fig. 22b | Two-way ANOVA | Sidak's test | BSA type: F (1, 8) = 10.32 | *P*=0.0124 |
| | | | Solution vs adsorbed: F (1, 8) = 88.47 | *P*<0.0001 |
| | | | Interaction: F (1, 8) = 2.579 | *P*=0.1470 |
| Supplementary Fig. 23c | Unpaired t-test | | t=1.503, df=4 | *P*=0.2074 |
| Supplementary Fig. 23d | Unpaired t-test | | t=3.599, df=4 | *P*=0.0228 |
| Supplementary Fig. 23g | Unpaired t-test | | t=1.612, df=4 | *P*=0.1823 |
| Supplementary Fig. 23h | Unpaired t-test | | t=4.188, df=4 | *P*=0.0138 |
| Supplementary Fig. 23k | Unpaired t-test | | t=1.247, df=4 | *P*=0.2803 |
| Supplementary Fig. 23l | Unpaired t-test | | t=2.393, df=4 | *P*=0.0749 |

**Supplementary Table 7. Statistical comparison of temperature-dependent BSA protein sizes from DLS measurements.**

Dunnett’s multiple comparisons test results after one-way ANOVA of the DLS-tracked size measurements of BSA proteins 1-6 as a function of temperature. Dunnett’s test was conducted for each BSA type, with the corresponding 25 °C data point as the reference point. Multiplicity-adjusted *P* values are reported.

| **Fig. 2a DLS BSA 1** | | | |
|---|---|---|---|
| Dunnett's multiple comparisons test | Significant? | Summary | Multiplicity-adjusted *P* value |
| BSA 1 25 °C vs 50 °C | No | ns | >0.9999 |
| BSA 1 25 °C vs 55 °C | No | ns | 0.9999 |
| BSA 1 25 °C vs 60 °C | No | ns | 0.997 |
| BSA 1 25 °C vs 65 °C | Yes | **** | <0.0001 |
| **Fig. 2a DLS BSA 2** | | | |
| BSA 2 25 °C vs 50 °C | No | ns | >0.9999 |
| BSA 2 25 °C vs 55 °C | No | ns | >0.9999 |
| BSA 2 25 °C vs 60 °C | No | ns | >0.9999 |
| BSA 2 25 °C vs 65 °C | No | ns | 0.9929 |
| BSA 2 25 °C vs 70 °C | Yes | **** | <0.0001 |
| **Fig. 2a DLS BSA 3** | | | |
| BSA 3 25 °C vs 50 °C | No | ns | >0.9999 |
| BSA 3 25 °C vs 55 °C | No | ns | >0.9999 |
| BSA 3 25 °C vs 60 °C | No | ns | >0.9999 |
| BSA 3 25 °C vs 65 °C | No | ns | >0.9999 |
| BSA 3 25 °C vs 70 °C | No | ns | >0.9999 |
| BSA 3 25 °C vs 75 °C | Yes | **** | <0.0001 |
| **Fig. 2a DLS BSA 4** | | | |
| BSA 4 25 °C vs 50 °C | No | ns | 0.9999 |
| BSA 4 25 °C vs 55 °C | No | ns | 0.9994 |
| BSA 4 25 °C vs 60 °C | No | ns | 0.5338 |
| BSA 4 25 °C vs 65 °C | Yes | **** | <0.0001 |
| **Fig. 2a DLS BSA 5** | | | |
| BSA 5 25 °C vs 50 °C | No | ns | >0.9999 |
| BSA 5 25 °C vs 55 °C | No | ns | 0.9998 |
| BSA 5 25 °C vs 60 °C | No | ns | 0.288 |
| BSA 5 25 °C vs 65 °C | Yes | **** | <0.0001 |
| **Fig. 2a DLS BSA 6** | | | |
| BSA 6 25 °C vs 50 °C | No | ns | >0.9999 |
| BSA 6 25 °C vs 55 °C | No | ns | 0.9998 |
| BSA 6 25 °C vs 60 °C | No | ns | 0.3531 |
| BSA 6 25 °C vs 65 °C | Yes | **** | <0.0001 |

**Supplementary Table 8. Statistical comparison of BSA protein sizes from DLS measurements conducted at 25 °C.**

Tukey’s multiple comparisons test results after one-way ANOVA comparing the DLS-tracked sizes of BSA proteins at 25 °C. Multiplicity-adjusted *P* values are reported.

| **Fig. 2a DLS BSA 1 – 6 at 25 °C** | | | |
|---|---|---|---|
| Tukey's multiple comparisons test | Significant? | Summary | Multiplicity-adjusted *P* value |
| BSA 1 vs. BSA 2 | No | ns | >0.9999 |
| BSA 1 vs. BSA 3 | No | ns | 0.9997 |
| BSA 1 vs. BSA 4 | No | ns | 0.9241 |
| BSA 1 vs. BSA 5 | No | ns | 0.9988 |
| BSA 1 vs. BSA 6 | No | ns | 0.9564 |
| BSA 2 vs. BSA 3 | No | ns | >0.9999 |
| BSA 2 vs. BSA 4 | No | ns | 0.8961 |
| BSA 2 vs. BSA 5 | No | ns | 0.9969 |
| BSA 2 vs. BSA 6 | No | ns | 0.9361 |
| BSA 3 vs. BSA 4 | No | ns | 0.8096 |
| BSA 3 vs. BSA 5 | No | ns | 0.9842 |
| BSA 3 vs. BSA 6 | No | ns | 0.8673 |
| BSA 4 vs. BSA 5 | No | ns | 0.9913 |
| BSA 4 vs. BSA 6 | No | ns | >0.9999 |
| BSA 5 vs. BSA 6 | No | ns | 0.9972 |

**Supplementary Table 9. Statistical comparison of QCM-D |$\Delta F_{max}$|, |$\Delta F_{max}/\Delta D_{max}$| values, and LSPR $\Delta\lambda_{max}$ values obtained from BSA adsorption measurements.**

Tukey's multiple comparisons test results after one-way ANOVA of the |$\Delta F_{max}$| and |$\Delta F_{max}/\Delta D_{max}$| values from QCM-D measurements and the $\Delta\lambda_{max}$ values from LSPR measurements for BSA proteins 1-6. Multiplicity-adjusted *P* values are reported.

| **Fig. 3d QCM-D \|$\Delta F_{max}$\|** | | | |
|---|---|---|---|
| Tukey's multiple comparisons test | Significant? | Summary | Multiplicity-adjusted *P* value |
| BSA 1 vs. BSA 2 | Yes | ** | 0.0012 |
| BSA 1 vs. BSA 3 | Yes | **** | <0.0001 |
| BSA 1 vs. BSA 4 | Yes | **** | <0.0001 |
| BSA 1 vs. BSA 5 | Yes | **** | <0.0001 |
| BSA 1 vs. BSA 6 | Yes | **** | <0.0001 |
| BSA 2 vs. BSA 3 | Yes | **** | <0.0001 |
| BSA 2 vs. BSA 4 | No | ns | 0.1595 |
| BSA 2 vs. BSA 5 | Yes | *** | 0.0003 |
| BSA 2 vs. BSA 6 | Yes | ** | 0.0018 |
| BSA 3 vs. BSA 4 | Yes | **** | <0.0001 |
| BSA 3 vs. BSA 5 | Yes | **** | <0.0001 |
| BSA 3 vs. BSA 6 | Yes | **** | <0.0001 |
| BSA 4 vs. BSA 5 | Yes | * | 0.0192 |
| BSA 4 vs. BSA 6 | No | ns | 0.1415 |
| BSA 5 vs. BSA 6 | No | ns | 0.8258 |
| **Fig. 3e QCM-D \|$\Delta F_{max}/\Delta D_{max}$\|** | | | |
| BSA 1 vs. BSA 2 | No | ns | 0.9996 |
| BSA 1 vs. BSA 3 | Yes | ** | 0.0063 |
| BSA 1 vs. BSA 4 | Yes | *** | 0.0007 |
| BSA 1 vs. BSA 5 | Yes | *** | 0.0004 |
| BSA 1 vs. BSA 6 | Yes | ** | 0.0055 |
| BSA 2 vs. BSA 3 | Yes | ** | 0.0038 |
| BSA 2 vs. BSA 4 | Yes | ** | 0.001 |
| BSA 2 vs. BSA 5 | Yes | *** | 0.0006 |
| BSA 2 vs. BSA 6 | Yes | ** | 0.0091 |
| BSA 3 vs. BSA 4 | Yes | **** | <0.0001 |
| BSA 3 vs. BSA 5 | Yes | **** | <0.0001 |
| BSA 3 vs. BSA 6 | Yes | **** | <0.0001 |
| BSA 4 vs. BSA 5 | No | ns | 0.9979 |
| BSA 4 vs. BSA 6 | No | ns | 0.7563 |
| BSA 5 vs. BSA 6 | No | ns | 0.5199 |
| **Fig. 3f LSPR $\Delta\lambda_{max}$** | | | |
| BSA 1 vs. BSA 2 | No | ns | 0.9724 |
| BSA 1 vs. BSA 3 | Yes | * | 0.0362 |
| BSA 1 vs. BSA 4 | Yes | * | 0.0413 |
| BSA 1 vs. BSA 5 | Yes | **** | <0.0001 |

| | | | |
|---|---|---|---|
| BSA 1 vs. BSA 6 | Yes | ** | 0.0012 |
| BSA 2 vs. BSA 3 | Yes | * | 0.0104 |
| BSA 2 vs. BSA 4 | No | ns | 0.1393 |
| BSA 2 vs. BSA 5 | Yes | *** | 0.0001 |
| BSA 2 vs. BSA 6 | Yes | ** | 0.004 |
| BSA 3 vs. BSA 4 | Yes | *** | 0.0002 |
| BSA 3 vs. BSA 5 | Yes | **** | <0.0001 |
| BSA 3 vs. BSA 6 | Yes | **** | <0.0001 |
| BSA 4 vs. BSA 5 | Yes | ** | 0.0091 |
| BSA 4 vs. BSA 6 | No | ns | 0.3279 |
| BSA 5 vs. BSA 6 | No | ns | 0.2951 |

**Supplementary Table 10. Statistical comparison of $(d\Delta\lambda/dt)_{max}$ values and ATR-FTIR helicity values obtained for BSA proteins 1-6.**

Tukey's multiple comparisons test results after one-way ANOVA of the $(d\Delta\lambda/dt)_{max}$ values and two-way ANOVA of helicity values of BSA proteins 1-6 from ATR-FTIR measurements. Helicity values from ATR-FTIR spectroscopy measurements are compared between BSA types in solution and in the adsorbed state, separately. Multiplicity-adjusted *P* values are reported.

| **Fig. 4a LSPR $(d\Delta\lambda/dt)_{max}$** | | | |
|---|---|---|---|
| Tukey's multiple comparisons test | Significant? | Summary | Multiplicity-adjusted *P* value |
| BSA 1 vs. BSA 2 | No | ns | 0.2397 |
| BSA 1 vs. BSA 3 | No | ns | 0.9383 |
| BSA 1 vs. BSA 4 | Yes | *** | 0.0001 |
| BSA 1 vs. BSA 5 | Yes | **** | <0.0001 |
| BSA 1 vs. BSA 6 | Yes | **** | <0.0001 |
| BSA 2 vs. BSA 3 | No | ns | 0.0575 |
| BSA 2 vs. BSA 4 | Yes | ** | 0.0041 |
| BSA 2 vs. BSA 5 | Yes | *** | 0.0002 |
| BSA 2 vs. BSA 6 | Yes | *** | 0.0009 |
| BSA 3 vs. BSA 4 | Yes | **** | <0.0001 |
| BSA 3 vs. BSA 5 | Yes | **** | <0.0001 |
| BSA 3 vs. BSA 6 | Yes | **** | <0.0001 |
| BSA 4 vs. BSA 5 | No | ns | 0.4657 |
| BSA 4 vs. BSA 6 | No | ns | 0.9221 |
| BSA 5 vs. BSA 6 | No | ns | 0.9383 |
| **Fig. 4b ATR-FTIR helicity (solution)** | | | |
| BSA 1 vs. BSA 2 | No | ns | 0.9543 |
| BSA 1 vs. BSA 3 | No | ns | 0.9543 |
| BSA 1 vs. BSA 4 | No | ns | 0.9061 |
| BSA 1 vs. BSA 5 | No | ns | 0.9543 |
| BSA 1 vs. BSA 6 | No | ns | 0.3489 |
| BSA 2 vs. BSA 3 | No | ns | >0.9999 |
| BSA 2 vs. BSA 4 | No | ns | 0.4417 |
| BSA 2 vs. BSA 5 | No | ns | 0.5433 |
| BSA 2 vs. BSA 6 | No | ns | 0.0759 |
| BSA 3 vs. BSA 4 | No | ns | 0.4417 |
| BSA 3 vs. BSA 5 | No | ns | 0.5433 |
| BSA 3 vs. BSA 6 | No | ns | 0.0759 |
| BSA 4 vs. BSA 5 | No | ns | >0.9999 |
| BSA 4 vs. BSA 6 | No | ns | 0.9061 |
| BSA 5 vs. BSA 6 | No | ns | 0.8364 |
| **Fig. 4b ATR-FTIR helicity (adsorbed)** | | | |
| BSA 1 vs. BSA 2 | No | ns | 0.9952 |
| BSA 1 vs. BSA 3 | No | ns | 0.9824 |
| BSA 1 vs. BSA 4 | Yes | * | 0.0169 |

| | | | |
|---|---|---|---|
| BSA 1 vs. BSA 5 | Yes | * | 0.025 |
| BSA 1 vs. BSA 6 | Yes | ** | 0.005 |
| BSA 2 vs. BSA 3 | No | ns | >0.9999 |
| BSA 2 vs. BSA 4 | Yes | ** | 0.005 |
| BSA 2 vs. BSA 5 | Yes | ** | 0.0076 |
| BSA 2 vs. BSA 6 | Yes | ** | 0.0014 |
| BSA 3 vs. BSA 4 | Yes | ** | 0.0033 |
| BSA 3 vs. BSA 5 | Yes | ** | 0.005 |
| BSA 3 vs. BSA 6 | Yes | *** | 0.0009 |
| BSA 4 vs. BSA 5 | No | ns | >0.9999 |
| BSA 4 vs. BSA 6 | No | ns | 0.9952 |
| BSA 5 vs. BSA 6 | No | ns | 0.9824 |

**Supplementary Table 11. Statistical comparison of the blocking efficiency of BSA proteins 1-6 against serum biofouling and to protect against silica nanoparticle-induced complement activation.**

Tukey's multiple comparisons test results after one-way ANOVA of the surface passivation performance evaluation of BSA proteins 1-6 to inhibit serum biofouling and to minimize nanoparticle-induced complement activation. Multiplicity-adjusted *P* values are reported.

| **Fig. 5c Serum biofouling blocking efficiency** | | | |
|---|---|---|---|
| Tukey's multiple comparisons test | Significant? | Summary | Multiplicity-adjusted *P* value |
| BSA 1 vs. BSA 2 | No | ns | 0.2275 |
| BSA 1 vs. BSA 3 | No | ns | 0.9852 |
| BSA 1 vs. BSA 4 | Yes | * | 0.0338 |
| BSA 1 vs. BSA 5 | Yes | *** | 0.0004 |
| BSA 1 vs. BSA 6 | Yes | *** | 0.0003 |
| BSA 2 vs. BSA 3 | No | ns | 0.0836 |
| BSA 2 vs. BSA 4 | No | ns | 0.8362 |
| BSA 2 vs. BSA 5 | Yes | * | 0.0174 |
| BSA 2 vs. BSA 6 | Yes | * | 0.0133 |
| BSA 3 vs. BSA 4 | Yes | * | 0.0115 |
| BSA 3 vs. BSA 5 | Yes | *** | 0.0002 |
| BSA 3 vs. BSA 6 | Yes | *** | 0.0001 |
| BSA 4 vs. BSA 5 | No | ns | 0.1245 |
| BSA 4 vs. BSA 6 | No | ns | 0.0967 |
| BSA 5 vs. BSA 6 | No | ns | >0.9999 |
| **Fig. 5d Protection against complement activation** | | | |
| BSA 1 vs. BSA 2 | No | ns | 0.3532 |
| BSA 1 vs. BSA 3 | No | ns | 0.7653 |
| BSA 1 vs. BSA 4 | Yes | * | 0.0244 |
| BSA 1 vs. BSA 5 | Yes | * | 0.0272 |
| BSA 1 vs. BSA 6 | No | ns | 0.0848 |
| BSA 2 vs. BSA 3 | No | ns | 0.9797 |
| BSA 2 vs. BSA 4 | Yes | *** | 0.0002 |
| BSA 2 vs. BSA 5 | Yes | *** | 0.0002 |
| BSA 2 vs. BSA 6 | Yes | *** | 0.0007 |
| BSA 3 vs. BSA 4 | Yes | *** | 0.001 |
| BSA 3 vs. BSA 5 | Yes | ** | 0.0011 |
| BSA 3 vs. BSA 6 | Yes | ** | 0.0041 |
| BSA 4 vs. BSA 5 | No | ns | >0.9999 |
| BSA 4 vs. BSA 6 | No | ns | 0.9917 |
| BSA 5 vs. BSA 6 | No | ns | 0.9945 |

**Supplementary Table 12. Statistical comparison of the adsorbed protein layer thickness of BSA proteins 1-6.**

Tukey's multiple comparisons test results after one-way ANOVA of the adsorbed protein layer thickness of BSA proteins 1-6 based on Voigt-Voinova modeling of QCM-D data. Multiplicity-adjusted *P* values are reported.

| **Supplementary Fig. 5 Effective thickness of adsorbed BSA layers** | | | |
|---|---|---|---|
| Tukey's multiple comparisons test | Significant? | Summary | Multiplicity-adjusted *P* value |
| BSA 1 vs. BSA 2 | Yes | **** | <0.0001 |
| BSA 1 vs. BSA 3 | Yes | ** | 0.0022 |
| BSA 1 vs. BSA 4 | Yes | **** | <0.0001 |
| BSA 1 vs. BSA 5 | Yes | **** | <0.0001 |
| BSA 1 vs. BSA 6 | Yes | **** | <0.0001 |
| BSA 2 vs. BSA 3 | Yes | **** | <0.0001 |
| BSA 2 vs. BSA 4 | No | ns | 0.9996 |
| BSA 2 vs. BSA 5 | Yes | ** | 0.0089 |
| BSA 2 vs. BSA 6 | No | ns | 0.0795 |
| BSA 3 vs. BSA 4 | Yes | **** | <0.0001 |
| BSA 3 vs. BSA 5 | Yes | **** | <0.0001 |
| BSA 3 vs. BSA 6 | Yes | **** | <0.0001 |
| BSA 4 vs. BSA 5 | Yes | ** | 0.0054 |
| BSA 4 vs. BSA 6 | Yes | * | 0.0487 |
| BSA 5 vs. BSA 6 | No | ns | 0.776 |

**Supplementary Table 13. Statistical comparison of the surface passivation performance of BSA proteins 1-6 to reduce silica nanoparticle-induced complement activation.**

Dunnett’s multiple comparisons test results after one-way ANOVA of the surface passivation performance of BSA proteins 1-6 to inhibit silica nanoparticle-induced complement activation. The Bare NP group served as the control. Multiplicity-adjusted *P* values are reported.

| **Supplementary Fig. 13 Normalized SC5b-9 levels** | | | |
|---|---|---|---|
| Dunnett's multiple comparisons test | Significant? | Summary | Multiplicity-adjusted *P* value |
| Bare NP vs. BSA 1 | Yes | **** | <0.0001 |
| Bare NP vs. BSA 2 | Yes | **** | <0.0001 |
| Bare NP vs. BSA 3 | Yes | **** | <0.0001 |
| Bare NP vs. BSA 4 | Yes | **** | <0.0001 |
| Bare NP vs. BSA 5 | Yes | **** | <0.0001 |
| Bare NP vs. BSA 6 | Yes | **** | <0.0001 |

**Supplementary Table 14. Statistical comparison of temperature-dependent CA-BSA 5 protein sizes from DLS measurements.**

Dunnett's multiple comparisons test results after one-way ANOVA of the DLS-tracked size measurements of CA-BSA 5 as a function of temperature. The 25 °C data point served as the reference point. Multiplicity-adjusted *P* values are reported.

| **Supplementary Fig. 17a DLS CA-BSA 5** | | | |
|---|---|---|---|
| Dunnett's multiple comparisons test | Significant? | Summary | Multiplicity-adjusted *P* value |
| CA-BSA 5 25 °C vs 50 °C | No | ns | >0.9999 |
| CA-BSA 5 25 °C vs 55 °C | No | ns | 0.9999 |
| CA-BSA 5 25 °C vs 60 °C | No | ns | >0.9999 |
| CA-BSA 5 25 °C vs 65 °C | No | ns | 0.9981 |
| CA-BSA 5 25 °C vs 70 °C | No | **** | <0.0001 |

**Supplementary Table 15. Statistical comparison of ATR-FTIR helicity values obtained from BSA 5 and CA-BSA 5 in solution and in the adsorbed state.**

Sidak's multiple comparisons test results after two-way ANOVA of helicity values of BSA 5 and CA-BSA 5 from ATR-FTIR measurements. Helicity values from ATR-FTIR spectroscopy measurements were compared between BSA types in solution and in the adsorbed state, separately. Multiplicity-adjusted *P* values are reported.

| **Supplementary Fig. 22b ATR-FTIR CA-BSA 5** | | | |
|---|---|---|---|
| Sidak's multiple comparisons test | Significant? | Summary | Multiplicity-adjusted *P* value |
| BSA 5 vs. CA-BSA 5 (solution) | No | ns | 0.4945 |
| BSA 5 vs. CA-BSA 5 (adsorbed) | Yes | * | 0.0185 |

### References


1. Schindelin, J. et al. Fiji: an open-source platform for biological-image analysis. *Nat. Methods* **9**, 676 (2012).
2. Arakawa, T. & Kita, Y. Stabilizing effects of caprylate and acetyltryptophanate on heat-induced aggregation of bovine serum albumin. *Biochim. Biophys. Acta, Protein Struct. Mol. Enzymol.* **1479**, 32-36 (2000).
3. Anraku, M. et al. Stabilizing mechanisms in commercial albumin preparations: octanoate and N-acetyl-L-tryptophanate protect human serum albumin against heat and oxidative stress. *Biochim. Biophys. Acta, Proteins Proteomics* **1702**, 9-17 (2004).
4. Mannuzza, F.J. & Montalto, J.G. Is Bovine Albumin Too Complex to Be Just a Commodity? *BioProcess Int.* (2010).
5. Boyer, P.D., Lum, F.G., Ballou, G.A., Luck, J.M. & Rice, R.G. The combination of fatty acids and related compounds with serum albumin I. Stabilization against heat denaturation. *J. Biol. Chem.* **162**, 181-198 (1946).
